# Global phylogeography and ancient evolution of the widespread human gut virus crAssphage

**DOI:** 10.1101/527796

**Authors:** Robert A. Edwards, Alejandro A. Vega, Holly M. Norman, Maria Ohaeri, Kyle Levi, Elizabeth A. Dinsdale, Ondrej Cinek, Ramy K. Aziz, Katelyn McNair, Jeremy J. Barr, Kyle Bibby, Stan JJ. Brouns, Adrian Cazares, Patrick A. de Jonge, Christelle Desnues, Samuel L. Díaz Muñoz, Peter C. Fineran, Alexander Kurilshikov, Rob Lavigne, Karla Mazankova, David T. McCarthy, Franklin L. Nobrega, Alejandro Reyes Muñoz, German Tapia, Nicole Trefault, Alexander V. Tyakht, Pablo Vinuesa, Jeroen Wagemans, Alexandra Zhernakova, Frank M. Aarestrup, Gunduz Ahmadov, Abeer Alassaf, Josefa Anton, Abigail Asangba, Emma Billings, Vito Adrian Cantu, Jane M. Carlton, Daniel Cazares, Gyu-Sung Cho, Tess Condeff, Pilar Cortés, Mike Cranfield, Daniel A. Cuevas, Rodrigo De la Iglesia, Przemyslaw Decewicz, Michael P. Doane, Nathaniel J. Dominy, Lukasz Dziewit, Bashir Mukhtar Elwasila, A. Murat Eren, Charles Franz, Jingyuan Fu, Cristina Garcia-Aljaro, Elodie Ghedin, Kristen M. Gulino, John M. Haggerty, Steven R. Head, Rene S. Hendriksen, Colin Hill, Heikki Hyöty, Elena N. Ilina, Mitchell T. Irwin, Thomas Jeffries, Juan Jofre Torroella, Randall E. Junge, Scott T. Kelley, Martin Kowalewski, Deepak Kumaresan, Steven Leigh, Eugenia S. Lisitsyna, Montserrat Llagostera, Julia M. Maritz, Linsey C. Marr, Angela McCann, Mohammadali Khan Mirzaei, Shahar Molshanski-Mor, Silvia Monteiro, Ben Moreira-Grez, Megan Morris, Lawrence Mugisha, Maite Muniesa, Horst Neve, Nam-phuong Nguyen, Olivia D. Nigro, Anders S. Nilsson, Taylor O’Connell, Rasha Odeh, Andrew Oliver, Mariana Piuri, Aaron J. Prussin II, Udi Qimron, Zhe-Xue Quan, Petra Rainetova, Adán Andrés Ramírez Rojas, Raul Raya, Gillian A.O. Rice, Alessandro Rossi, Ricardo Santos, John Shimashita, Elyse N. Stachler, Lars C. Stene, Ronan Strain, Rebecca Stumpf, Pedro J. Torres, Alan Twaddle, MaryAnn Ugochi Ibekwe, Nicolás Villagra, Stephen Wandro, Bryan White, Andy Whitely, Katrine L. Whiteson, Cisca Wijmenga, Maria M. Zambrano, Henrike Zschach, Bas E. Dutilh

## Abstract

**Abstract:** Microbiomes are vast communities of microbes and viruses that populate all natural ecosystems. Viruses have been considered the most variable component of microbiomes, as supported by virome surveys and examples of high genomic mosaicism. However, recent evidence suggests that the human gut virome is remarkably stable compared to other environments. Here we investigate the origin, evolution, and epidemiology of crAssphage, a widespread human gut virus. Through a global collaboratory, we obtained DNA sequences of crAssphage from over one-third of the world’s countries, and showed that its phylogeography is locally clustered within countries, cities, and individuals. We also found colinear crAssphage-like genomes in both Old-World and New-World primates, challenging genomic mosaicism and suggesting that the association of crAssphage with primates may be millions of years old. We conclude that crAssphage is a benign globetrotter virus that may have co-evolved with the human lineage and an integral part of the normal human gut virome.

## Main text

Phages form the vast majority of the human gut virome in healthy individuals, with an estimated ~5 x 10^9^ phages per gram of human feces versus ~9 x 10^10^ bacteria^1,2^. Phages are critical for the control of bacterial populations and vary widely between individuals^3–5^. Evolutionary and genomic studies have suggested that dynamic phage-host interactions are reflected in phage genomes, which show high sequence diversity and mosaicism^6,7^. In marine aquatic ecosystems phages only persist in the environment for one to two days^8–10^, but those dynamics may be drastically different in the human gut microbiome, where phages can persist for over a year^3,11^.

### crAssphage populations in the human gut

We assessed the origin, evolution, and epidemiology of crAssphage, one of the most ubiquitous human gut viruses to understand the stability of the human gut virome. We previously recovered the crAssphage sequence from over half of 466 fecal metagenomics datasets^12^ and the first cultured member of the expansive crAss-like family^13^ was recently reported^14^. We screened the crAssphage genome for regions that were present in many different datasets, where variable segments were flanked by conserved regions suitable for targeting by PCR primers, identifying three amplicon regions of ~1.3 kilobases (see Methods). We tested fecal samples from 45 healthy individuals from four cities on two continents and found that almost half of these volunteers (21 individuals) were crAss-positive as determined by gel electrophoresis. We followed six individuals over two months to assess the stability of crAssphage populations (Fig. 1). Two individuals were consistently crAss-negative, while others displayed more variable dynamics. Notably, DNA sequencing revealed that crAssphage strains from each individual tended to be phylogenetically clustered, although the sequences are not phylogenetically clustered by date (Fig. 1G-I). This suggests that multiple, closely related crAssphage populations may co-exist within one individual whose abundance, and thus whose detection varies in time. Such ecological dynamics may also explain the fact that Male 1 was intermittently crAss-negative, although his sequences are still clustered in the phylogenetic trees.

**Fig. 1.**
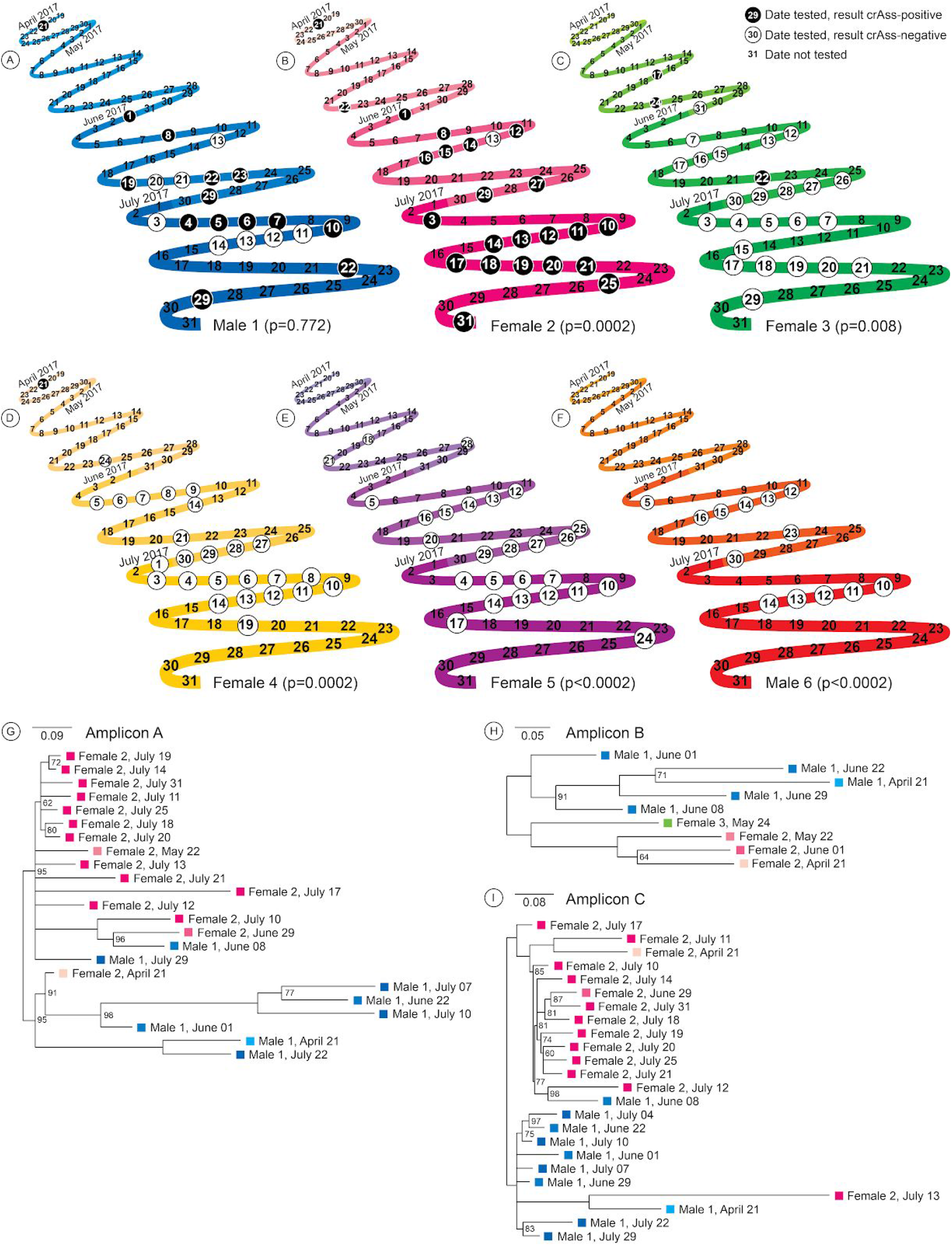
CrAssphage presence/absence status is stable over time in the human gut. A-F: Timelines showing the crAssphage status of six volunteers between April and July 2017, where every zig or zag represents a week from Monday to Sunday and subsequent months are indicated in increasingly intense colors per individual. On the circled dates, individuals were PCR tested for crAssphage using amplicons A, B, and C, and gel electrophoresis of the three amplicons was always consistent per sample. Black and white circles indicate crAss-positive and crAss-negative samples, respectively. G-I: Unrooted maximum likelihood phylogenies of amplicons A-C show clustering of the sequences by volunteer (note: not all crAss-positive samples could be sequenced). Branches with <60% bootstrap support were collapsed, values <100% are displayed. As in panels A-F, colors correspond to the individual and the month in which the sample was taken.

To confirm the intra-individual evolution observed above, we recovered twenty different crAssphage genomes from the fecal viromes of three adult female twin pairs and their mothers, using the same datasets that we originally used to discover crAssphage^3,12^ and built a phylogenomic tree. Genomes sampled up to one year apart from the same individual clustered together in the tree (Extended Data Fig. 1), consistent with a model of intra-individual evolution of these dominant gut virome populations that are generally acquired once, but may diverge into several different, but related subpopulations with time^3,11^.

### CrAssphage is globally distributed and locally clustered

The phylogenies in Fig. 1G-I and Extended Data Fig. 1 suggested that individuals have a dominant and stable crAssphage population in their gut microbiome, but these results might be skewed by PCR amplification or metagenome assembly. While higher order groups including species and genera remain controversial in viral taxonomy and depend on complete genome sequences^15,16^, strains can readily be defined as unique sequences^17^. To analyze how many strains could co-occur within one sample, we downloaded 95,552 metagenomics datasets from all environments from the Sequence Read Archive^18^. Using a strain-resolved bioinformatics pipeline developed for this analysis^19^ (see Methods), we extracted the three amplicon regions from 2,216 datasets, most of which contained only a single crAssphage strain (Fig. 2). While 95% of all recovered strains were only found in a single sample, one strain of amplicon C was identified 104 times in different datasets (listed in Supplementary File 1), showing the exceptional ubiquity of some strains around the world. It has been suggested that crAssphage is not acquired early in life^20^, but our global analysis identified crAssphage in at least 134 infant samples (26 with locality information, see Supplementary File 2), confirming recent incidental findings that crAssphage can be found in infants^20,21^. Sixteen metagenomes contain more than 100 strains. Phylogenetic trees containing these sequences showed that, as in Figure 1 and Extended Data Figure 1, strains within a single individual tend to be recently diverged, although different co-occurring clusters could be observed in some cases (Extended Data Fig. 2). Interestingly, the two samples with the most diverse crAssphage populations are from young individuals, including a healthy USA child^22^ containing up to 1,409 strains and a one-year old Finnish infant^23^ containing up to 748 strains (Fig. 2; Supplementary File 3). Still, our phylogenomic tree based on the twin study^3^ suggests that crAssphage is not always vertically transmitted since none of the daughter strains cluster with their mothers (Extended Data Fig. 1).

**Fig. 2.**
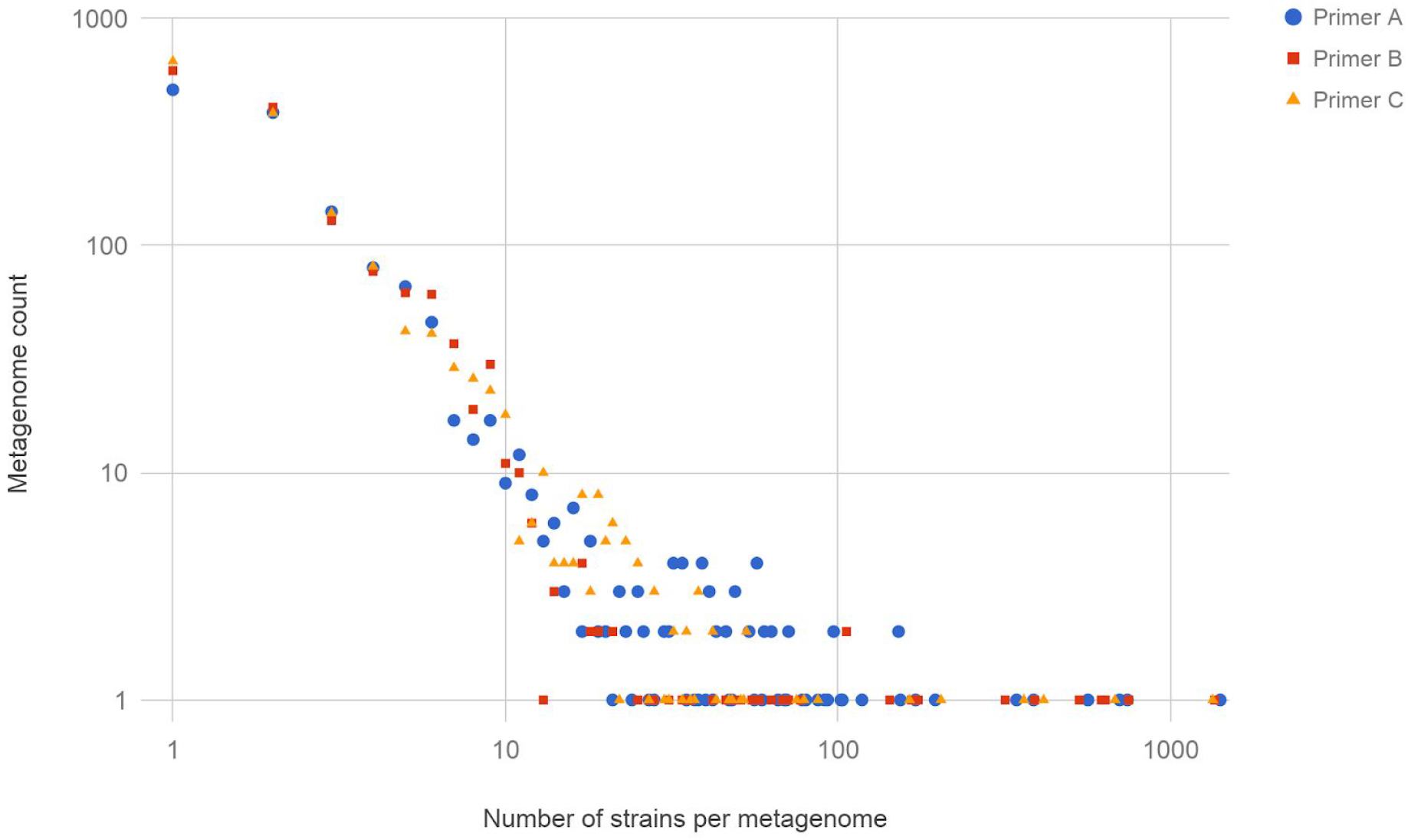
Diversity of crAssphage strains in metagenomic samples. Strains for three amplicon regions A, B, and C were detected with Gretel^19^ in 2,216 metagenomes (see Supplementary File 3).

To investigate the global phylogeography of crAssphage, we collected data about the three amplicon regions from various sources and combined them in a large-scale phylogenetic analysis, providing the first worldwide overview of the evolution of an epitome of the human gut virome (Extended Data Table 1). First, we launched a global collaboratory to amplify and sequence the three regions of the crAssphage genome from local sites. To obtain the highest expected rate of detection, collaborators sampled wastewater treatment plants. We combined these sequences with data from the COMPARE sewage sampling project (http://www.compare-europe.eu/), and the sequences from our metagenomics searches and individual volunteers found above. Together, we analyzed 32,273 different crAssphage sequences from at least 67 countries on six continents (34% of the countries in the world, see Fig. 3, Extended Data Fig. 3, and Supplementary File 2). We reconstructed phylogenetic trees for the subset of strains with locality information and assessed the distribution of associated sampling metadata by using permutation statistics^24^. Sequences from the same country, location, and sampling date are significantly clustered in the phylogeny (*p*<0.001, see Extended Data Fig. 4), and the genetically most similar other strain tends to be geographically close (Fig. 3). Thus, crAssphage is a cosmopolitan inhabitant of the human gut the world-over, with a geographically and temporally local sequence signature that may prove useful in future forensic applications of fecal contamination identification and detection^25–28^.

**Fig. 3.**
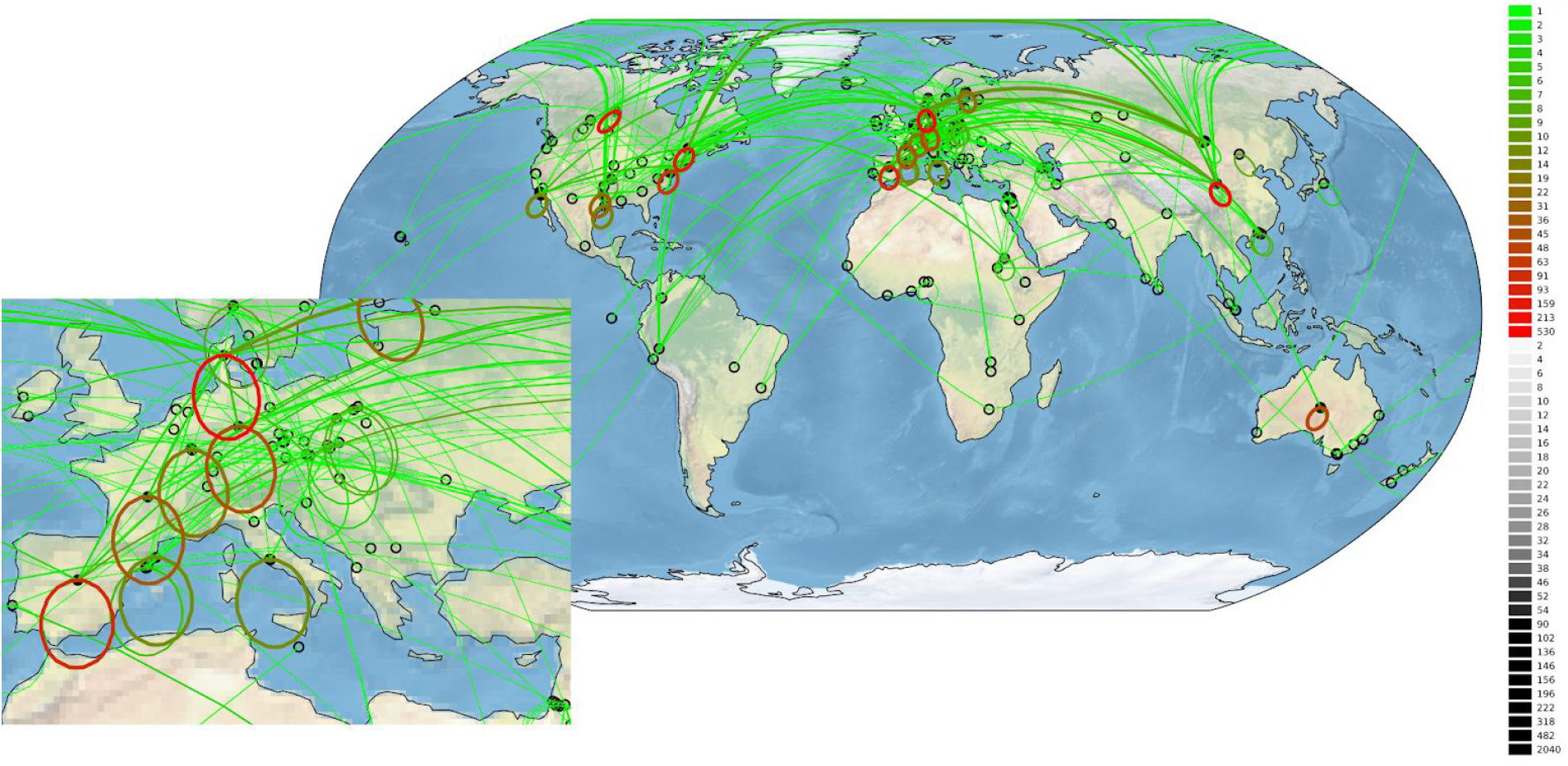
Global locations of 2,424 crAssphage strains (amplicon A, see Extended Data Fig. 3 for amplicons B and C). The number of strains at each location is reflected in the intensity of the black circles. For each strain, a link to the genetically most similar other strain is indicated with a line, geographically very close connections (<100 km) are indicated with circled lines for visibility. The color of the lines indicates the number links between two locations. Inset: samples from Europe.

### CrAssphage has evolved with humans

The global distribution of crAssphage led us to ask the question whether this virus was present in early humans and has evolved with us as we spread out and colonized the planet. Alternatively, and consistent with the view of viruses as rapidly evolving entities, it is possible that crAssphage emerged recently, perhaps through recombination of other viruses, and spread around the world either via factors related to the human host, e.g. the global food supply chain or international travel, or via the epidemiology of our intestinal bacteria.

To assess the possible ancient association of crAssphage-like phages with the human lineage, we screened the datasets from our global data survey for remote human populations. We found a few crAssphage-like sequences in fecal samples from rural Malawi and from the Amazonas of Venezuela^29^ (see Extended Data Table 2). In contrast, mummified gut samples from three pre-Columbian Andean mummies^30^ and the European iceman^31^ were all crAss-negative. While this could suggest that these individuals were crAss-negative, it is also possible that any crAssphage DNA has degraded over thousands of years.

Next, we sequenced and assembled fifteen fecal metagenomes from five species of non-human primates to search for crAssphage in our distant primate relatives. None of the assembled nucleotide sequences matched the amplicon regions used above, as only short stretches of nucleotide homology were identified to the crAssphage genome^32^. Surprisingly, many short homologous regions were found in several long sequences of ~90,000 nucleotides, and when displayed in a dot-plot, revealed a range of near-complete, distant crAssphage relatives in apes, Old-World monkeys, and New-World monkeys (Fig. 4). While those genomes were distantly related to crAssphage they were clearly colinear, showing the long-term genomic stability of this widespread gut virus. These results are consistent with a recent study that identified ten candidate crAss-like phage genera, whose genomes were also colinear^33^. To investigate the phylogenetic relationships between those genomes and the ones identified in the non-human primates, we created a concatenated alignment phylogeny of 15 proteins. As shown in Extended Data Fig. 5, the non-human primate sequences are related to candidate genera III and IX, two candidate genera of the Alphacrassvirinae to which the prototypical crAssphage candidate genus I also belongs^33^. While most non-human primate sequences form deep clades, two sequences from Gorilla_1 are closely related to the human strains CDZH01002743 (25 year old male from Canada^34^) and FDYN_MS_11 (healthy individual from Ireland^33^). We hypothesize that this strain may have been transmitted from humans, as this gorilla has has human contact (http://gracegorillas.org/2017/12/29/pinga/). She also contains an additional strain that clusters among the other non-human primate sequences in candidate genus IX.

**Fig. 4.**
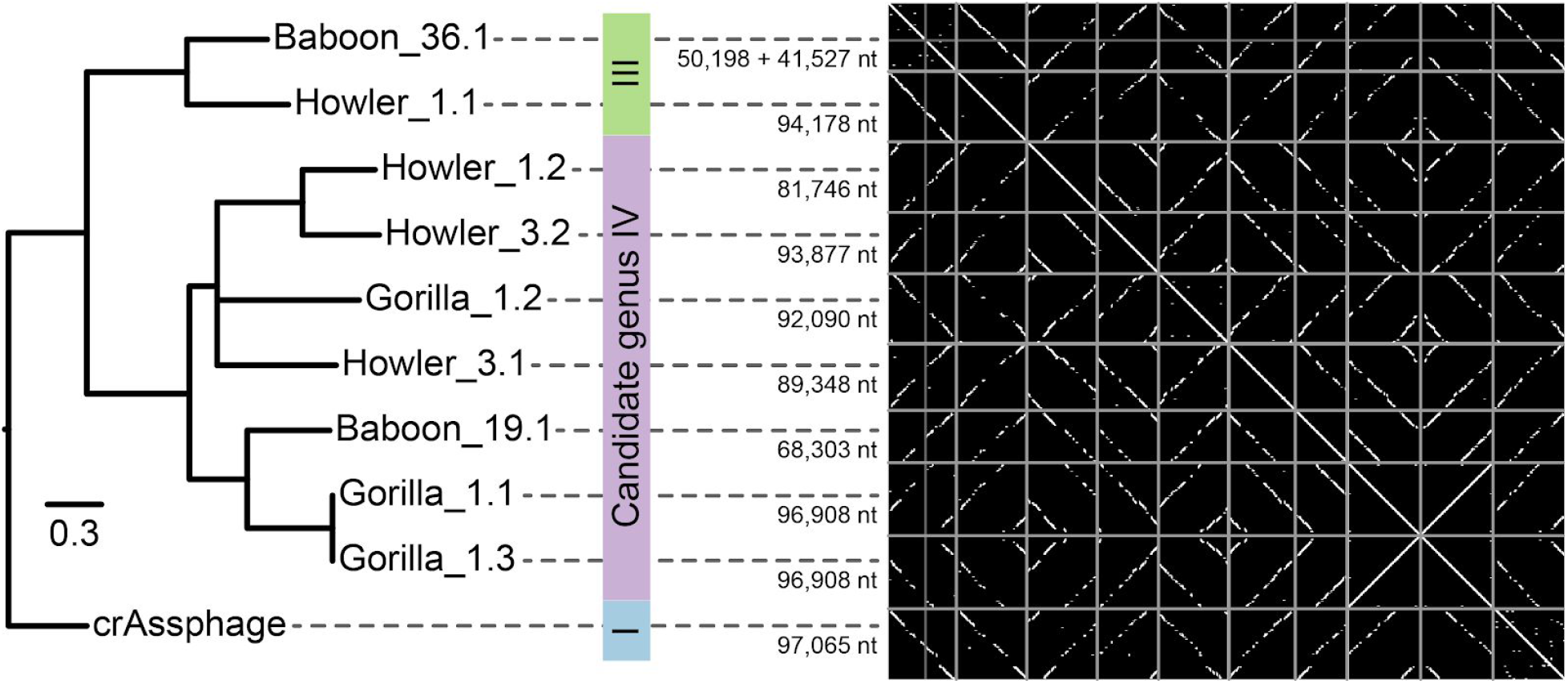
Maximum likelihood phylogeny and dotplot showing full genomic colinearity between crAssphage and ten long contigs assembled from fecal metagenomes of different non-human primates. The phylogeny is based on a concatenated trimmed protein alignment of 15 homologous ORFs. The tree is rooted as suggested in^33^ and candidate genera are indicated with colored blocks. All branches had 100% bootstrap support, one exception <50% was collapsed. The scale bar indicates 0.3 mutations per site. For a phylogeny of all 119 Alphacrassvirinae and non-human primates see Extended Data Fig. 5. Dotplots are based on high-scoring segment pairs (blastn E-value <0.001) between all contigs. The figure is to scale, numbers to the left of the dotplot indicating genome or contig lengths. Note that circular permutation of some genomes leads to apparently broken diagonals in some dotplots.

### CrAssphage belongs to the normal human virome

To investigate the association of crAssphage with characteristics of the human host and its microbiome, we investigated the correlation between fecal crAssphage abundance and a range of host factors and microbial taxa. By exploiting shotgun metagenomes and host metadata from the LifeLines-DEEP cohort^36,37^, we correlated the abundance of crAssphage across 1,135 individuals with 207 exogenous and intrinsic human variables, including 78 dietary factors, 41 intrinsic factors, 39 diseases, 44 drug groups, 5 smoking categories (Supplementary File 4), and 490 microbial taxa (Supplementary File 5). We found significant but weak correlations with several diet categories (Benjamini-Hochberg <5% false discovery rate), including protein, carbohydrates, and caloric intake, basic food groups that are probably related to the dietary preferences of the crAssphage host bacteria^12,36,38–40^. The most significant correlations of crAssphage with microbial taxa in the LifeLines-DEEP cohort included the family *Prevotellaceae*, consistent with our previous prediction that crAssphage infects bacteria of the *Bacteroidetes* phylum^12^. Diverse dietary associations have been observed for different *Bacteroidetes* members, including the genus *Bacteroides* that was linked to a long-term Western diet rich in animal protein and sugars^41^, while *Prevotella* and *Paraprevotella* were linked to low protein and high fiber^42^. The most reliable computational phage-host signal to date^43^ is a 100% matching CRISPR spacer in *Porphyromonas* sp. 31_2 isolated from human feces^13^, another species within the *Bacteroidetes*, and the first cultured crAss-like phage was recently isolated by using *Bacteroides intestinalis* as an isolation host^14^. Given the potentially family-scale taxonomic diversity of crAssphages^13^, it is likely that they infect a range of hosts throughout the *Bacteroidetes* phylum, leading to poor abundance correlations between crAssphage and specific host taxa. The LifeLines-DEEP cohort did not reveal a significant relationship between crAssphage and any human health or disease parameters, consistent with a previous study showing absence of an association with diarrhea^21^. As crAssphage abundance is not related to any health-related variables, we conclude that it is a part of the normal human virome^44^.

## Conclusions

The human gut virome mainly consists of phages that infect the abundant and diverse bacteria living in our gut. Phages are generally thought of as transient entities in the environment, whose fast infection cycle and relatively error-prone replication machinery enable rapid co-evolution with their hosts, which in turn would be reflected in highly diverse viral (meta-)genome sequences^6,7^. Indeed, we found thousands of crAssphage strains throughout human feces-associated environments around the world. These strains are geographically and temporally clustered, consistent with rapid evolution and local dispersion. However, we also identified one exceptionally widespread strain in up to 104 different samples from e.g. Denmark, France, Germany, Israel, Italy, Japan, and USA (Supplementary File 1). We suggest that this conservation primarily reflects recent spread by human global migration, although a crAssphage strain with potentially high fitness or environmental stability cannot be ruled out. Moreover, we identified highly divergent but fully colinear genome sequences from the crAss-like candidate genera III and IX^33^ in all major groups of primates, suggesting that crAssphage has had a stable genome structure for millions of years, and a stable association with the primate lineage and its microbiome^45^ since our early ancestors began their great migration out of Africa.

Recently, the extent of gene flux and genomic mosaicism has been proposed to differ between temperate and virulent phages^46^. Virulent phages tend to be genomically stable, while temperate phages fall into either high or low gene flux modes. Our results challenge high genomic mosaicism in viruses, showing that phage genome structure can be remarkably conserved in the stable environment provided by the human gut. Based on our observations, components of the human gut virome may be remarkably stable over millions of years, possibly reflecting the environmental stability of its niche. This high stability of the primate gut also limits the ability of its specialized microbes and viruses to escape to other environments. Indeed, this specificity makes crAssphage one of the strongest human fecal contamination markers to date^25–27^. Taken together, our results provide the first global overview of the phylogeography of one of the most abundant and widespread viruses in the human gut, with evidence of both an ancient evolution and ongoing local dispersion.

## Supporting information

Most ubiquitous crAssphage strain

Global Sampling of crAssphage

Gretel strains

Lifelines phenotype correlations

Lifelines microbial correlations

## Acknowledgments

- Special thanks to the following individuals who provided access to wastewater treatment samples: Robert Matthews, Mitchell Wright, John Alexander, Susie Arredondo, Nicki Branch, Doug Campbell, Ravy Chea, Dawn McDougle, Jeff Parks, and Vasana Vipatapat.
- We thank the Mountain Gorilla Veterinary Project, and the Maryland Zoo for collecting the gorilla fecal samples in Rwanda. We thank Gillian Britton for collecting the baboon fecal samples in Ethiopia. We thank the Chimpanzee Sanctuary and Wildlife Conservation Trust CSWCT), the Uganda Wildlife Authority (UWA), and the Uganda National Council for Science and Technology (UNCST) for collecting the chimpanzee fecal samples in Uganda.
- Primate samples were provided by the Primate Microbiome Collaborative (PMC) at the University of Illinois Urbana Champaign. Gorilla samples were originally obtained by MK and the Mountain Gorilla Veterinary Project in Rwanda. GB and ND provided the wild baboon samples from Ethiopia. Howler samples were provided by MK and lemur samples were provided by RJ and MI. RMS and LM provided the chimpanzee samples with permission from the Chimpanzee Sanctuary and Wildlife Conservation Trust CSWCT), the Uganda Wildlife Authority (UWA), and the Uganda National Council for Science and Technology (UNCST).
- We thank Joseph Manor at Central Virology Laboratory, Chaim Sheba Medical Center, Tel-Hashomer Hospital for help with sample collection.
- The primate microbiome project was supported by NSF BCS 0935347 to SL, RMS, BW, KN.
- We thank the COMPARE and LifeLines-DEEP projects for sharing data.
- ODN thanks Dr. Grieg Steward, University of Hawai’i, Manoa for support.
- PCF thanks Dr. Corinda Taylor for support with the PCR.

## Funding

The funders had no role in study design, data collection and analysis, decision to publish, or preparation of the manuscript.

- This work used the Extreme Science and Engineering Discovery Environment (XSEDE) Jetstream resources at Indiana University and Texas Advanced Computing Center through allocation MCB170036 to RAE, which is supported by National Science Foundation grant number ACI-1548562.
- Some of this work was supported by San Diego State University Grants Programs to RAE including the Summer Undergraduate Research Program.
- This work was supported by National Science Foundation grant numbers MCB-1441985 to RAE and DUE-1323809 to EAD.
- This work was supported by the Department of Energy Lawrence Livermore National Laboratory grant B618146 to RAE.
- PAJ and BED were supported by the Netherlands Organization for Scientific Research (NWO) Vidi grant 864.14.004.
- FLN was supported by the Netherlands Organization for Scientific Research (NWO) Veni grant 016.Veni.181.092.
- SJJB was supported by European Research Council Stg grant [638707] and the Netherlands Organization for Scientific Research (NWO) Vidi grant 864.11.005.
- OC and KM were supported by Ministry of Health of the Czech Republic grants nr. 15-31426A and 15-29078A.
- PCF was supported by a Rutherford Discovery Fellowship from the Royal Society of New Zealand.
- JJB was supported by the Australian Research Council (ARC) Discovery Early Career Researcher Award (DE170100525).
- SLDM was supported by an NIH Pathway to Independence Fellowship (1K99AI119401-01A1).
- DTM thanks the Australian Research Council’s Linkage Project LP160100408, Melbourne Water and EPA Victoria for funding the collection of samples in Melbourne.
- KB was supported by award number 1510925 from the United States National Science Foundation.
- MTI was supported by National Geographic Society (CRE) and NSERC.
- CD was supported by Agence Nationale de la Recherche JCJC grant #ANR-13-JSV6-0004 and Investissements d’Avenir Méditerranée Infection #10-IAHU-03.
- The LifeLines-DEEP sample collection and analysis was funded by the Netherlands Heart Foundation (IN-CONTROL CVON grant 2012-03 to AZ and JF), by the Top Institute Food and Nutrition, Wageningen, the Netherlands (TiFN GH001to CW), by the Netherlands Organization for Scientific Research (NWO) Vidi grant 864.13.013 to JF, NWO Vidi grant 016.178.056 to AZ, NWO Spinoza Prize SPI 92-266 to CW), and by the European Research Council (ERC) FP7/2007-2013/ERC Advanced Grant agreement 2012-322698 to CW, ERC Starting Grant 715772 to AZ. AZ also holds a Rosalind Franklin Fellowship from the University of Groningen.
- The COMPARE data collection was funded by The Novo Nordisk Foundation (NNF16OC0021856).

## Author Contributions

BED, RAE conceived of the study, performed the experiments and bioinformatics, and wrote the paper with input from all authors. AAV performed the volunteer experiments and sampled San Diego wastewater treatment plants. FLN, HMN, MO, PAJ performed human volunteer experiments. AE, AR, AVT, DAC, JMH, KL, KMcN, TOC, VAC performed bioinformatics analysis. AARR, AAl, ACz, AMcC, AO, ARM, ASN, AW, BM, BME, CD, CF, CH, DC, DK, DTM, EAD, EB, ENI, ENS, ESL, GA, GCA, GSC, GT, HH, HN, JAB, JJB, JJT, JMC, JMM, JW, KB, KLW, KM, LCS, LD, MAUI, MKM, ML, MMZ, MMo, MMu, MP, MPD, NT, NV, OC, ODN, PC, PCF, PD, PR, PV, RI, RKA, RL, RO, RR, RSa, RSr, SJJB, SLDM, SM, SMM, SW, TC, TJ, UQ, ZXQ performed sampling, PCR, and sequencing. AK, AZ, CW, JF performed the Lifelines analysis. FMA, HZ, RSH provided and analysed COMPARE project data. AAs, BW, GAOR, NJD, NPN, RSu, SL analyzed and provided the non human primate sequences. MC collected Gorilla samples. AT, EG, KMG performed the NYC sewage sampling and data analysis. AJP, JS, LCM, PJT, SRH, STK examined crAssphage transfer among infants. MTI, REJ collected lemur sample. MK collected Howler monkey samples. LM collected chimpanzee samples.

## Methods

### Phylogenomic tree of twin study crAssphages

To assess the evolution of the intra-individual crAssphage population and its within-family relations, we assembled twenty different, near-complete crAssphage genomes from the fecal viromes of three female twin pairs and their mothers^1^ using SPAdes 3.11.0 with default metagenomics settings^2^. Contigs related to crAssphage were identified by querying the contigs against the crAssphage reference genome sequence (RefSeq identifier NC_024711.1) with blastn^3^ 2.5.0+ (E-value <0.001). Next, the bitscore (which is independent of the database size) was summed for each SPAdes contig, and contigs with a total summed bitscore ≥4,000 were selected. Note that shorter contigs with homology to crAssphage existed in the datasets, but we limited our analysis to the longest contigs with the strongest similarity signal to the crAssphage genome.

A phylogenomic tree (Extended Data Fig. 1) was created based on the near-complete crAssphage genomes from the twin gut viromes. First, open reading frames (ORFs) were identified on all contigs with Prodigal v2.6.3^4^ and queried against the crAssphage genome with blastp^3^ 2.5.0+ (E-value <0.001). Proteins were excluded that were missing from more than three genomes. This resulted in a dataset of 68 proteins that were aligned using Clustal Omega^5^ 1.2.0 with default parameters (crAssphage proteins orf00003, orf00007, orf00009, orf00010, orf00011, orf00012, orf00013, orf00014, orf00015, orf00016, orf00017, orf00018, orf00020, orf00022, orf00023, orf00024, orf00025, orf00026, orf00027, orf00029, orf00031, orf00032, orf00033, orf00035, orf00037, orf00038, orf00040, orf00041, orf00042, orf00044, orf00045, orf00046, orf00047, orf00053, orf00054, orf00055, orf00056, orf00057, orf00059, orf00060, orf00062, orf00063, orf00065, orf00066, orf00067, orf00068, orf00070, orf00071, orf00072, orf00074, orf00075, orf00078, orf00079, orf00080, orf00081, orf00082, orf00084, orf00086, orf00088, orf00091, orf00092, orf00093, orf00094, orf00095, orf00096, orf00097, orf00098, and orf00099). The aligned proteins were concatenated to form a superalignment of 25,066 residues, that was converted to an approximate maximum likelihood tree with IQ-tree^6,7^ 1.5.5 (options -alrt 1000 -bb 1000), which was shown to be the most robust phylogenetic method^8,9^.

### PCR primer design

PCR primers were designed to facilitate the identification of crAssphage in diverse sampling sites around the world, and amplify a variable region of the genome for phylogenetic analysis (Extended Data Tables 3-5). Several studies designed primers for selected crAssphage proteins^10–14^ but we took a data-driven approach by identifying regions of the crAssphage genome suitable for phylogeographic analysis, i.e. variable regions that were flanked by conserved regions that might be targeted by the primers. We identified these regions by determining the consensus sequence of the full crAssphage genome in 148 datasets where at least 10,000 reads were aligned to crAssphage in our previous study^15^. We used Bowtie2^16^ to map metagenomic sequencing reads against the crAssphage reference genome, and called the consensus using Samtools^17^, yielding 148 aligned consensus sequences. Next, we analyzed the genome for suitable regions according to the following criteria: 1) high diversity flanked by conserved regions; 2) present in ≥90% of all sequences (<10% gaps); 3) length between 1,000-1,400 nucleotides. From the resulting candidate regions, we identified potential PCR primer sites with a melting temperature between 70-72°C for further analysis. We defined three primer regions, that we call A, B, and C, and that amplify the regions primer A: 25634 .. 25653 bp; primer B: 33709 .. 33728 bp; and primer C: 43820 .. 43841bp in the canonical crAssphage genome that has RefSeq ID NC_024711.1 ^15^.

### PCR amplification and sequencing of primer regions

The metagenomics-guided primer design outlined above yielded 11 promising primer regions of the crAssphage genome, and following testing using raw sewage influent from four sewage plants in Southern California (see below), a standard protocol was developed for three regions of the crAssphage genome. To prepare the DNA template, the raw sewage influent (the stuff coming into the sewage plant) was centrifuged briefly to remove the solids, and passed through a 0.2 μm or 0.22 μm filter. 7 μl of supernatant was used in a 50 μl PCR reaction (Extended Data Table 4) with thirty cycles of amplification (Extended Data Table 5).

CrAss-status was determined by the identification of a band via gel electrophoresis, and sequencing was performed by Sanger sequencing using commercial providers. Bases were identified from the .ab1 files using phred^18,19^ version 0.071220.b, and overlapping reads were merged using merger from the EMBOSS suite (version 6.5.7.0) with default alignment parameters^20^. Sequences were then formatted so that the sequence identifier contained metadata about the sequences. Specifically, we recorded the collection location address, latitude, longitude, country, and altitude; the date of collection, the source of the sample (raw sewage, feces, etc), and other notes about the sample (see Supplementary File 2).

### Global crAssphage collaboratory

We initiated a global and local survey of crAssphage using an open science collaboratory framework. Scientists were asked to donate their time, expertise, and resources to collect samples from local sewage treatment plants, PCR amplify three regions of the crAssphage genome, and sequence those regions. To avoid potential contamination from usage of central reagent stocks, each lab was responsible for ordering their own primers, performing the PCR, and sequencing the end products. Primers were only provided to researchers involved in the project in a few cases, usually where ordering primers was too financially onerous. For those cases the primers were ordered from Integrated DNA Technologies (Coralville, IA) and provided to the researchers before the tubes had been opened. Notably, the sequences resulting from those cases did not show any clustering in our resulting analyses, ruling out the possibility of cross-contamination.

Our global survey of crAssphage showed the first direct evidence of a globally distributed phage associated with humans and wastewater treatment plants. We generated 544 crAssphage sequences (184 amplicon A; 158 amplicon B; 202 amplicon C) from 70 different locations in 23 countries across five continents. In most cases, a single pure sequence was obtained from each PCR amplification. This either suggests that there is a single dominant crAssphage strain in the environment, or that the PCR amplification resulted in one genotype being dominantly amplified at the expense of other sequences.

We tested for PCR amplification bias in two different ways. First, we started with three sewage samples from different wastewater treatment plants. From each sample we extracted five aliquots and amplified each in a separate reaction. All fifteen products were sequenced, and we recovered identical DNA sequences within wastewater plants but not between wastewater plants. Second, 30 different PCR fragments were cloned into the vector pTZ57R/T (Thermo Scientific) and sequenced independently (see below). Some sewage samples yielded mixed populations of sequences, while other sewage samples generated identical sequences.

Two wastewater treatment sites (Greymouth, New Zealand and Leuven, Belgium) identified mixed samples (e.g. Extended Data Fig. 6). In these cases, the sequences were identical at the beginning but abruptly degraded, possibly caused by amplification of two different genotypes that differ by a small insertion or deletion. Such an indel would place the sequence out of register, prohibiting the resolution of a single sequence from the trace data.

Only a few sites were unable to amplify crAssphage from the sewage using any of the primers. It is not clear whether that was due to a lack of crAssphage DNA in the sample, or potential contaminants in the sample that inhibited the PCR reaction. We deliberately did not provide a positive control for crAssphage to avoid cross contamination of samples. This is the first time a single phage genome has been sampled across almost the entire globe, and demonstrates the ubiquitous spread of this phage. Thus, while care was taken to provide the same protocol to all collaborators and crAssphage was identified at many sites, we focus only on positives in this study because negatives could still represent a problem with the experiments rather than lack of the phage in the sample.

### Sampling of volunteers

The volunteer sampling was conducted under IRB Approved Protocol Number HS-2016-0056 and BUA Protocol 17-02-003E from San Diego State University. In San Diego, we tested twelve American and one British individuals, eleven from San Diego and two from Irvine, between April 21 and May 25^th^, 2017; 4/7 males and 2/6 females were crAss-positive, as determined by PCR and gel-electrophoresis. Three crAss-positive and three crAss-negative volunteers were followed weekly until July 31^st^ 2017. On two separate weeks each volunteer was followed daily. Each volunteer was provided with swubes (Becton Dickenson, Franklin Lakes, NJ) to collect fecal samples immediately after defecation. Samples were processed by adding approximately 0.5 ml of sterile phosphate-buffered saline to the swube and placing the suspended material in an eppendorf tube. DNA extraction using the QIAamp PowerFecal DNA Kit (Qiagen, Hilden, Germany) was performed according to the manufacturer’s protocol. Samples were tested by PCR and all PCR products were sequenced by Eton Bioscience, San Diego. Another 32 volunteers from Wageningen and Utrecht were tested, where amplicon C was used for initial detection and amplicon B for confirmation. 15 people were crAss-positive (11 male, 4 female) and 17 were crAss-negative (14 male, 3 female). There were, 1 American (crAss-negative), 1 Australian (crAss-negative), 1 Chinese (crAss-negative), 1 Colombian (crAss-positive), 23 Dutch (11 crAss-positive, 12 crAss-negative), 1 German (crAss-positive), 1 Greek (crAss-positive), 1 Portuguese (crAss-positive), and 2 Spanish (crAss-negative). Interestingly, at least one couple living together for several years had a discordant crAss-status (data not shown).

### COMPARE global sewage sampling

Samples from 81 sewage plants in 63 countries were taken within the COMPARE project (http://www.compare-europe.eu/) for strain resolved metagenomic sequencing. Samples were spun down and DNA was isolated after the DNA isolation QIAamp Fast DNA Stool protocol^21^. Sequencing was performed at the Oklahoma Medical Research Foundation where the DNA was sheared to ~300 bp and library preparation was done using the NEXTflex PCR-free DNA-seq library preparation kit. The multiplexed samples were sequenced on a HiSeq3000 using 2×150 bp paired end. Subsequently, data was quality trimmed and assembled with SPAdes 3.9.0 using the -meta flag. CrAssphage contigs were identified as explained below for the primates’ fecal metagenomes. Amplicon regions were identified by using blastn 2.5.0+, and hits included if they overlapped >50% with the amplicon regions. Note that 158 out of 179 hits overlapped ≥99% with the amplicon regions. All E-values were equal to 0.0 in this small database consisting only of the COMPARE crAssphage contigs.

### Strain-resolved metagenomics

The sequence read archive^22^ (SRA) contains approximately ten petabases of DNA sequence (10^16^ bp), including data from many metagenomes. We developed a new pipeline to search the SRA using the Jetstream platform^23–25^. Initially we screened the 95,552 metagenomes identified by PARTIE^26^ for the presence of crAssphage by comparing 100,000 reads from each metagenome to the crAssphage reference genome sequence using bowtie2^16^. Metagenomes that had one or more matching reads in this initial screen were compared to the crAssphage genome to identify any sequencing reads in those metagenome libraries that match crAssphage. All metagenomic sequences were cleaned using our new parallel version of Prinseq, called Prinseq++ (https://github.com/Adrian-Cantu/PRINSEQ-plus-plus^27,28^). Sequences were trimmed to ensure the mean quality score was at least 20, no N’s were included in the sequences, all sequences were dereplicated, the ends were trimmed based on quality score cutoff, and each read was required to be a minimum 30 nucleotides long. The sequences were mapped and indexed using Bowtie2^16^ to generate a bam file. Details of the screening procedure are provided at https://github.com/linsalrob/SearchSRA^25^. A total of 10,260 metagenomes had matches to crAssphage over 1kb (Extended Data Fig. 7A). The variability of the three amplicon regions is shown in Extended Data Fig. 7B-D.

Entire amplicon regions were recovered from 2,216 metagenomes derived from 121 SRA Bioprojects, and those were used as input to the Gretel^29^ for probabilistic haplotype recovery. Initially, SNPs were identified from the bamfiles using snpper from Gretel-Test (https://github.com/SamStudio8/gretel-test) for each of the three regions used in the PCR. Variants predicted by Gretel were combined into a single fasta file for downstream analysis^29^. As in the global collaboratory, we focused on the crAssphage positive samples only. Due to persistent inconsistencies in the metadata of metagenomes submitted to SRA, we avoided an extensive search for e.g. all human fecal samples and/or all sewage or wastewater samples. Instead, we identified crAssphage in all metagenomic datasets, regardless of their environmental origin, and in our study refrain from making statements about the percentage of crAss-positive individuals based on this analysis. We observed a weak (r^2^ = 0.66) but statistically significant correlation (*p* < 0.01) between the depth of coverage of the three amplicon regions in the metagenomes and the number of strains recovered from each of the three amplicon regions (Extended Data Fig. 8). This may be expected because additional rare variants may be detected with deeper sequencing.

### Global phylogenetic trees of three amplicon regions

Using the methods outlined above, we amassed sequence data for each of the three amplicon regions from several different sources (Extended Data Fig. 8, Extended Data Table 1, and Supplementary File 2). As shown in Extended Data Table 1, only a subset of the sequences contained locality information, and could thus be included in the global phylogeographic analysis.

All sequences were then processed through the same pipeline that is provided as a *Makefile*^30^ in the GitHub repository (https://github.com/linsalrob/crassphage). The trees can be built using the GNU Make program. After alignment with MUSCLE^31^ version 3.8.31 (using a maximum of two iterations and with the -diags option to find diagonals), alignments were trimmed to remove any columns in the alignment that had gaps in more than 10% of the sequences by using a custom written Python program, which caused some of the sequences to be deleted (see Extended Data Fig. 3). Phylogenetic trees were constructed with IQ-tree^6^ (default settings) with ModelFinder^7^. The MUSCLE alignments and IQ-tree analysis were performed on a 540 node compute cluster in the Edwards Bioinformatics Lab. Trees were visualized with iTOL^32^.

### Assessment of metadata clustering

We assessed geographical clustering of crAssphage in the phylogeny, and clustering by sampling date. To obtain meaningful statistics, we developed a permutation approach as in^33^ that kept the branching structure of the phylogeny intact and reshuffled the leaf labels *N* times, each time asking if the geography and sampling date were more clustered in the randomly permuted than in the original tree. To account for phylogenetic noise in the tree topology, we increasingly collapsed branches with low bootstrap values. The statistics for the global phylogenies are presented in Extended Data Fig. 4 and empirical *p*-values for the clustering were calculated as explained below, resulting in *p*<0.001 for all statistics at all bootstrap levels.

To assess the extent of geographical clustering of crAssphage in the global phylogeny, we measured three different statistics. (1) For each branch in the tree we measured the frequency of the most frequently annotated country or locality, and averaged across all branches to get one “consistency” clustering statistic for the whole tree. (2) We counted the number of branches where all leaves have the same country or locality annotation, yielding a “perfect branches” statistic. (3) We calculated the standard deviation of all the pairwise geographical distances between leaves in a branch based on latitude/longitude coordinates, and averaged across all branches of the tree.

To assess phylogenetic clustering of the sampling dates, we calculated the standard deviation of all dates within a branch in the eight-digit numerical format YYYYMMDD, and averaged across all branches of the tree. Moreover, we also calculated the “consistency” and “perfect branches” statistics as above, by treating each date as a categorical rather than a numerical value.

### Rural Malawi and Amazonas of Venezuela

To investigate the presence of crAssphage in the fecal microbiota of human populations that were relatively remote from Western culture, we used metagenomic sequencing data from people from rural Malawi and the Amazonas of Venezuela^34^. Datasets were retrieved from MG-RAST^35^ and compared to the crAssphage reference genome sequence using Bowtie2^16^. As shown in Extended Data Table 6, a few reads mapped from both the Malawi and Venezuelan samples. As these hits did not cover the amplicon regions sampled in our global analysis, the sequences were not included in the global phylogeny.

### Mummies

To investigate the presence of crAssphage in the mummified fecal remains of ancient humans, we used metagenomic sequencing data from three pre-Columbian Andean mummies^36^ and the 5,300-year-old intestinal content of the tyrollean glacier mummy, Ötzi^37^. For the three pre-Columbian Andean mummies, sequences were downloaded from MG-RAST^35^ (MG-RAST project identifier mgp13354; 12 samples, 11,517,4154 reads, and 11,488,857,080 bp). For Ötzi^37^, sequences were downloaded from the SRA (ENA project identifier ERP012908; 43 samples, 2,797,498,968 reads, and 282,547,395,768 bp). Datasets were compared to the crAssphage reference genome sequence using bowtie2^16^ (nucleotide search) and RAPsearch2^38^ with E-value threshold <10^-5^ (protein search). No hits were found.

### Candidate crAss-like genera

Recently, Guerin *et al*. identified ten proposed crAssphage genera by reconstructing genomes from metagenomes^39^. They identified 249 genomes in total, and ascribed 63 genomes to candidate genus I, the genus that contains the prototypical crAssphage. Each of those 63 genomes contained the three amplicon regions described here, as detected using blastn, while none of the 186 genomes that belonged to other candidate genera contained any sequence similarity to those regions. Thus we are only identifying here are members of candidate genus I that infect bacteria in the human intestine.

### Primates

Fecal samples were collected from five species of primates in their natural habitats or in rehabilitation and conservation centres. Three baboon samples were collected from wild specimens (old-world monkeys). Baboon 19 and Baboon 22 are *Papio hamadryas/Papio anubis* hybrids from Awash, Ethiopia and Baboon 36 is a *Papio hamadryas* from Filwoha, Ethiopia. Three black and gold howler monkey samples were collected from wild *Alouatta caraya* in Argentina (new-world monkeys). Three lemur (sifakas) samples were collected from wild *Propithecus diadema* in Tsinjoarivo, Madagascar. Three eastern lowland gorilla *(Gorilla beringei graueri)* samples were taken at the Mountain Gorilla Veterinary Project. The apes were originally wild from Rwanda but were cared for in the Mountain Gorilla sanctuary when samples were taken. They have thus had some close contact with humans. Three chimpanzee samples were collected from *Pan troglodytes schweinfurthii* in Ngamba Island, Uganda. These apes were also sanctuary animals rescued from Congo and Uganda and have relatively close contact with humans. Metagenomic DNA libraries were constructed with the TruSeq DNA Sample Prep kit (Illumina, San Diego, CA, USA). All of the sequencing was done at the Roy J. Carver Biotechnology Center’s High-Throughput Sequencing and Genotyping Unit at the University of Illinois, Urbana-Champaign.

Metagenomic sequences were assembled, and contigs related to crAssphage identified as for the crAssphage genomes from the twin study (above). Shorter contigs with homology to crAssphage existed in the primate fecal metagenomic datasets, but we limited our analysis to the longest contigs with the strongest similarity signal to the crAssphage genome (total summed blastn bitscore ≥4,000) to validate the existence of ancient relatives of crAssphage in primate feces. The ten selected primate contigs contained a strong colinearity signal with the crAssphage genome, suggesting that they were near-complete genomes. Two contigs from Baboon 36 were merged, because they shared 66 overlapping nucleotides at the end of the contigs, had very similar assembly depth (7.7x and 7.4x, respectively), and were homologous to two non-overlapping sections of the crAssphage genome. Gorilla 1.1 and Gorilla 1.3 represent two nearly identical strains (one polymorphism in 96,908 nucleotides) that were independently recovered from the same ape, showing the robustness of the metagenomic sequencing and assembly approach. The sequences from Baboon 36 and Howler 1 were most similar to candidate genera III described by Guerin *et al*., while the other sequences were most similar to candidate genera IX, measured by the fraction of the genomes aligned with blastn and as shown in Fig. 4.

After identifying ORFs in all contigs with Phanotate v1.0.1^40^, homologous groups were identified by querying against proteins from the crAssphage genome with blastp 2.7.1+ (E-value <=10^-5^), the 15 protein homologs that were identified from crAssphage, the non-human primate, and the Alphacrassvirinae were separately aligned with MAFFT^41^ version 7.407, and concatenated. Positions with gaps in >5% of the sequences, and positions where every amino acid was identical, were removed. A maximum likelihood phylogenomic tree was created based on concatenated protein alignment with IQ-tree^6^ version 1.5.5 with ModelFinder^7^, tree search, 1,000 ultrafast bootstraps, and a SH-aLRT test (i.e. the IQ-tree options -alrt 1000 -bb 1000).

### Correlations with host factors and intestinal microbes

To explore the association between crAssphage abundance in the gut and a broad range of exogenous and intrinsic human phenotypes, as well as intestinal microbial taxa, we used data from the LifeLines-DEEP study^42^. The LifeLines-DEEP cohort is population-representative cohort of the citizens of northern Netherlands comprised of 1,135 individuals. Methods of sample collection, DNA extraction and sequencing, and phenotype selection were previously described^42^. To estimate the abundance of crAssphage in the LifeLines-DEEP samples, metagenomic sequencing data was mapped to the reference crAssphage genome using BWA version 0.7.15-r1140 with default parameters. CrAssphage relative abundance was calculated as the number of mapped reads divided by the total number of reads in the sample. Next, we used the Spearman rank sum test to estimate the association between crAssphage abundance and the phenotypes or microbial taxa of interest. Adjustment for multiple testing was conducted using the Benjamini-Hochberg procedure^43^. The results are listed in Supplementary Files 4-5.

### Data Availability

Sequence data that support the findings of this study have been deposited in GenBank under BioProject accession PRJNA510571. Each of the samples has a unique BioSample Accession number from SAMN10656826 through SAMN10658627 and SAMN10658653 and SAMN10659294.

In addition, all data, code, and analysis are available on GitHub under the MIT license. The data and code may be found at https://github.com/linsalrob/crAssphage. The current release is version 2.0 and has DOI: 10.5281/zenodo.1230436 ^44^.

## Extended Data Figures

**Extended Data Fig. 1.**
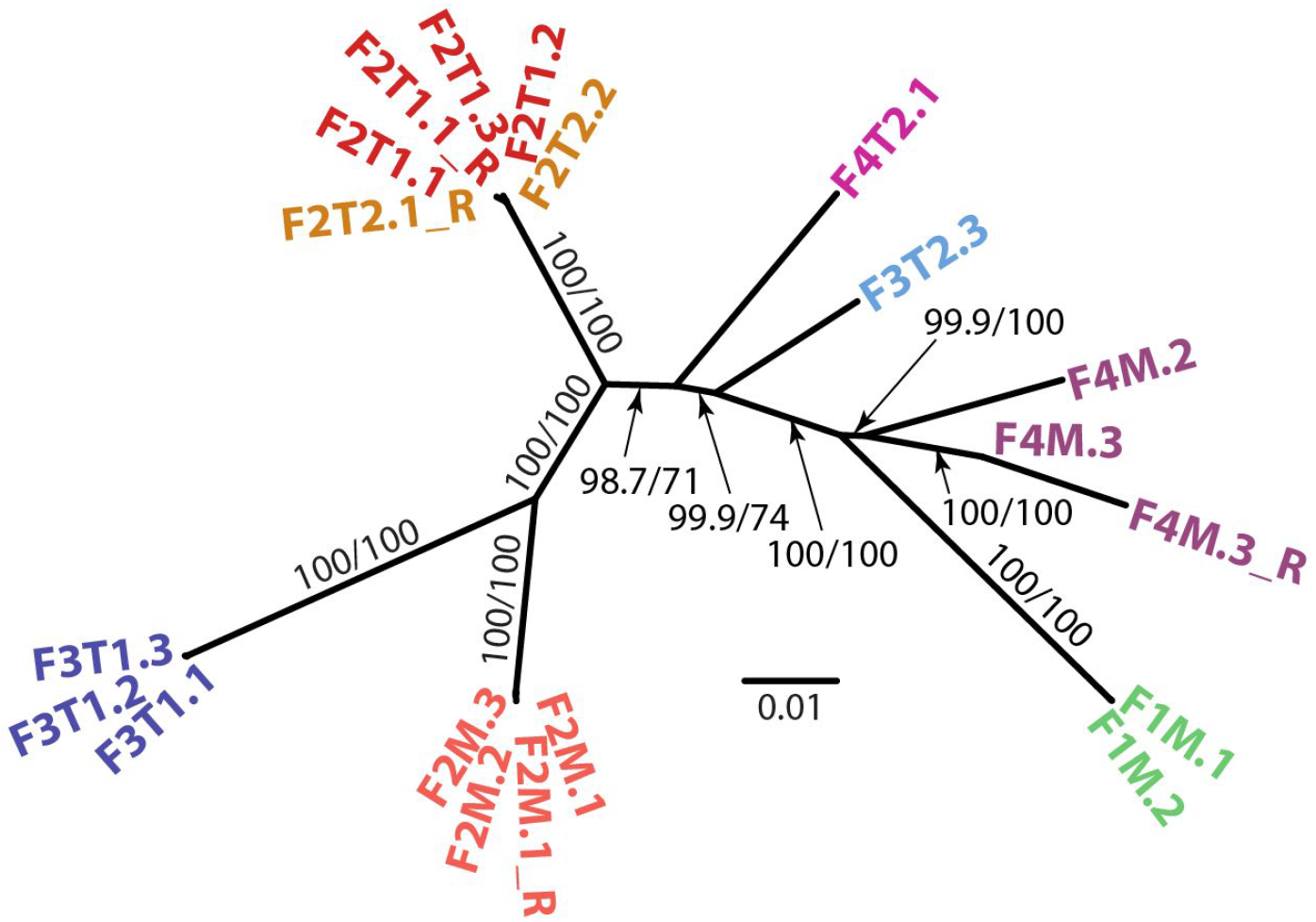
Phylogenomic tree of crAssphage genome sequences assembled from the Reyes twin study shows clustering of the strains by individual, with some samples taken up to one year apart^1^ yet clustering together in the tree. Sample tags indicate the family number (F1 through F4) and mother (M) or twins (T1 and T2). We could not reconstruct complete crAssphage genomes from F1T1, F1T2, F3M, and F4T1. Branch support values are SH-aLRT and ultrafast bootstrap support^2^. The scale bar indicates 0.01 mutations per site of the concatenated protein alignment.

**Extended Data Fig. 2.**
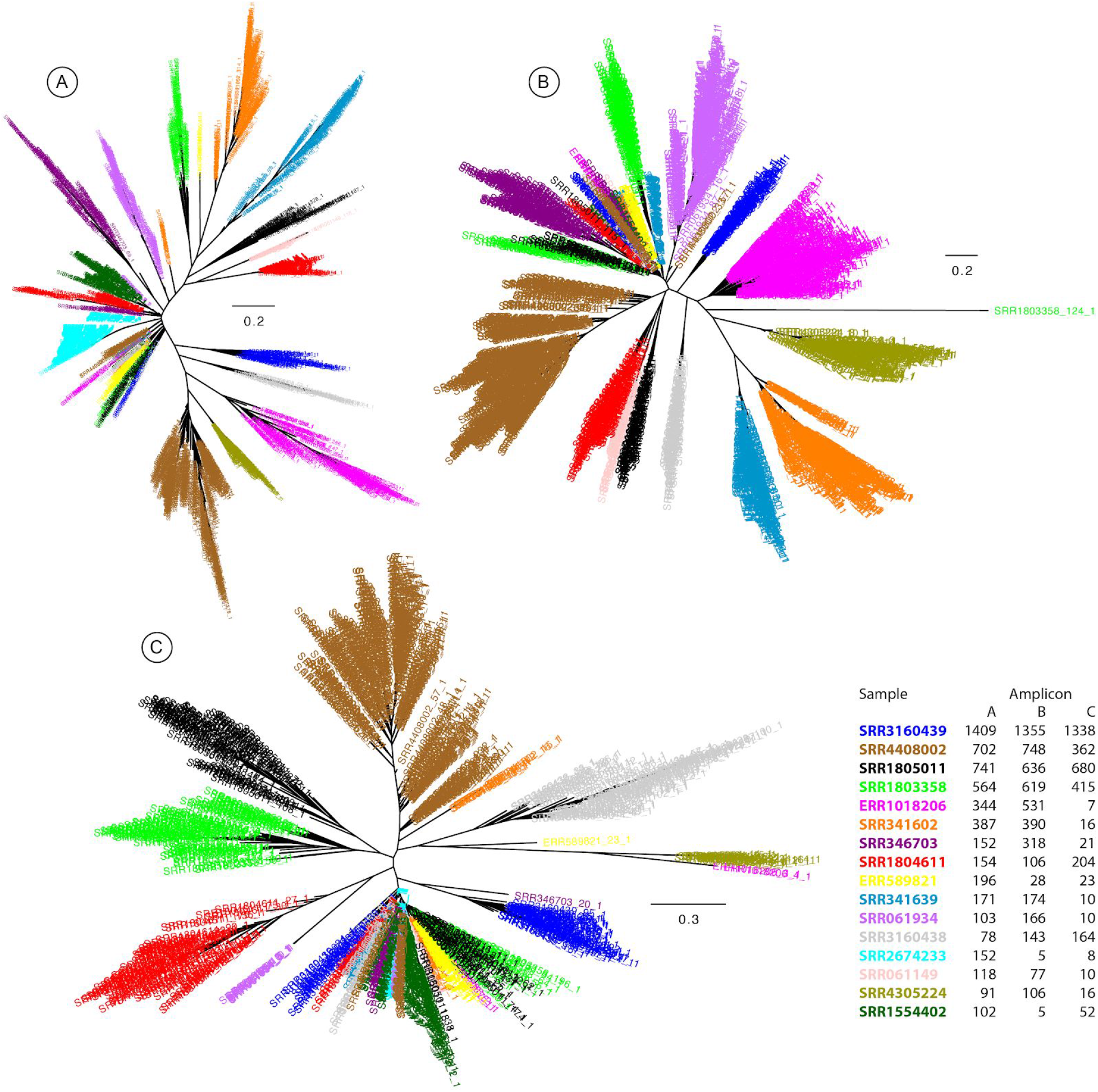
Phylogenetic trees of the three amplicons (A, B, and C) from the sixteen samples with more than 100 strains identified for any of the amplicons (see Supplementary File 3). The leaves of the trees are colored by sample, showing the strong phylogenetic relatedness of co-occurring crAssphage strains. The table in the inset lists the number of strains identified in each sample for each amplicon.

**Extended Data Fig. 3.**
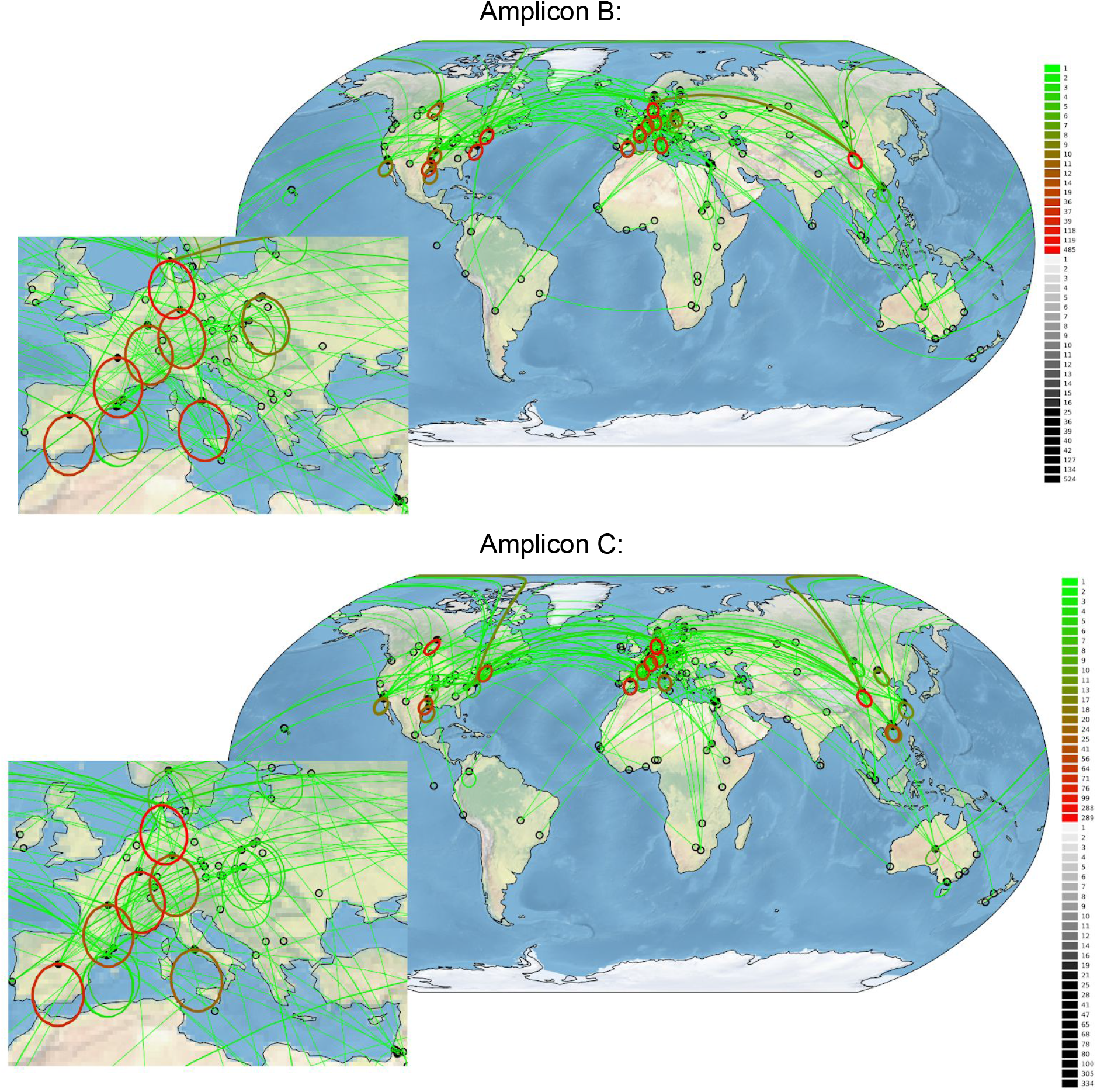
Global locations of 1,896 and 1,774 sequences from amplicons B and C, respectively. The number of strains at each location is reflected in the intensity of the black circles. For each strain, a link to the genetically most similar other strain is indicated with a line, geographically very close connections (<100 km) are indicated with circled lines for visibility. The color of the lines indicates the number links between two locations. Inset: samples from Europe.

**Extended Data Fig. 4.**
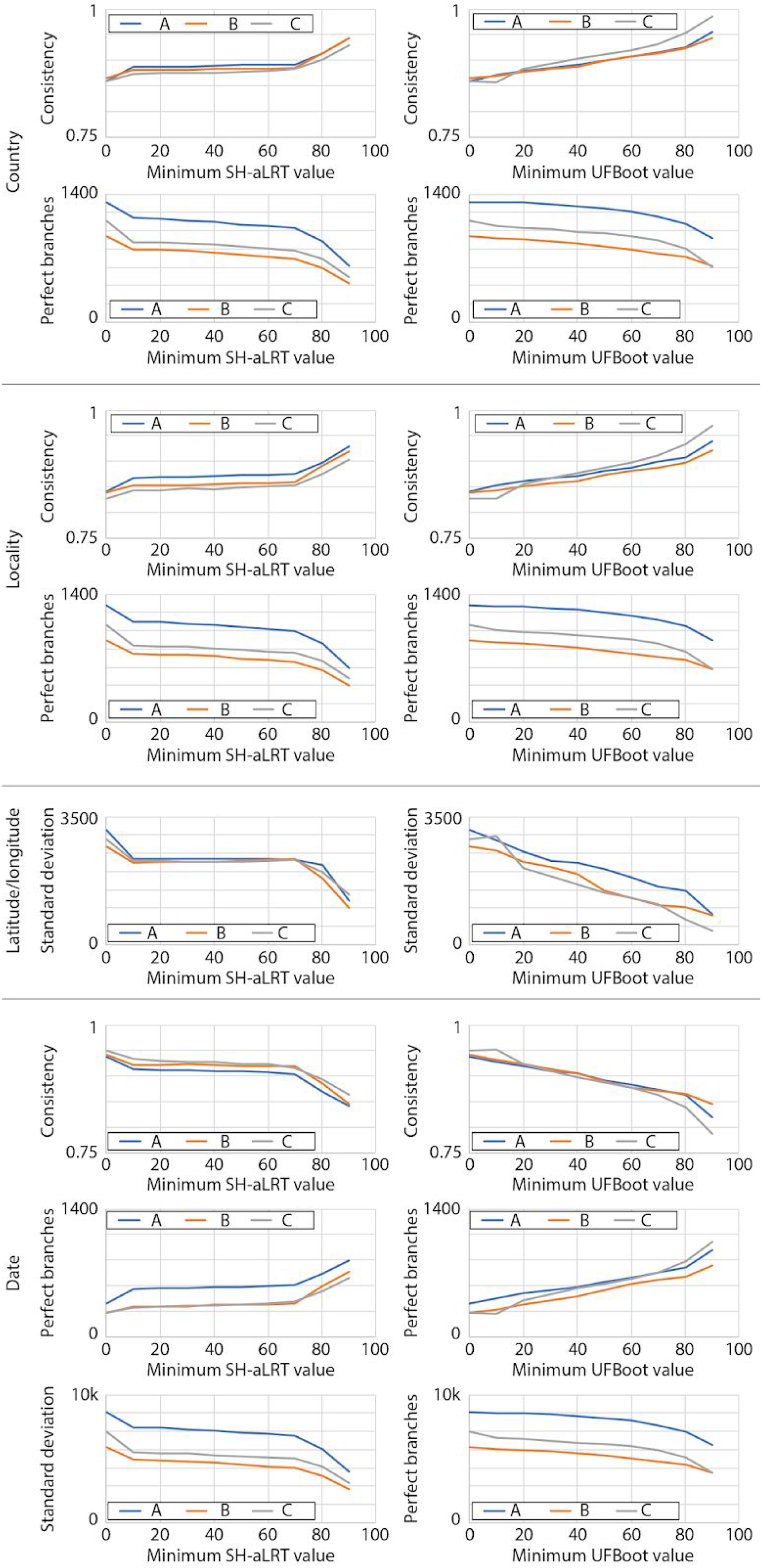
Geographical and temporal clustering statistics in the global phylogenetic trees of amplicon regions A (1,900 leaves), B (1,368 leaves), and C (1,621 leaves). Branches with increasing bootstrap values were collapsed (IQ-tree provides SH-aLRT and UFBoot bootstrap values, see left and right panels, respectively) and the statistics calculated. Next, statistics were also calculated based on 1,000 permutations of the leaf labels in the phylogenetic tree, but these statistics were never higher than with the original leaf labels so p<0.001.

**Extended Data Fig. 5.**
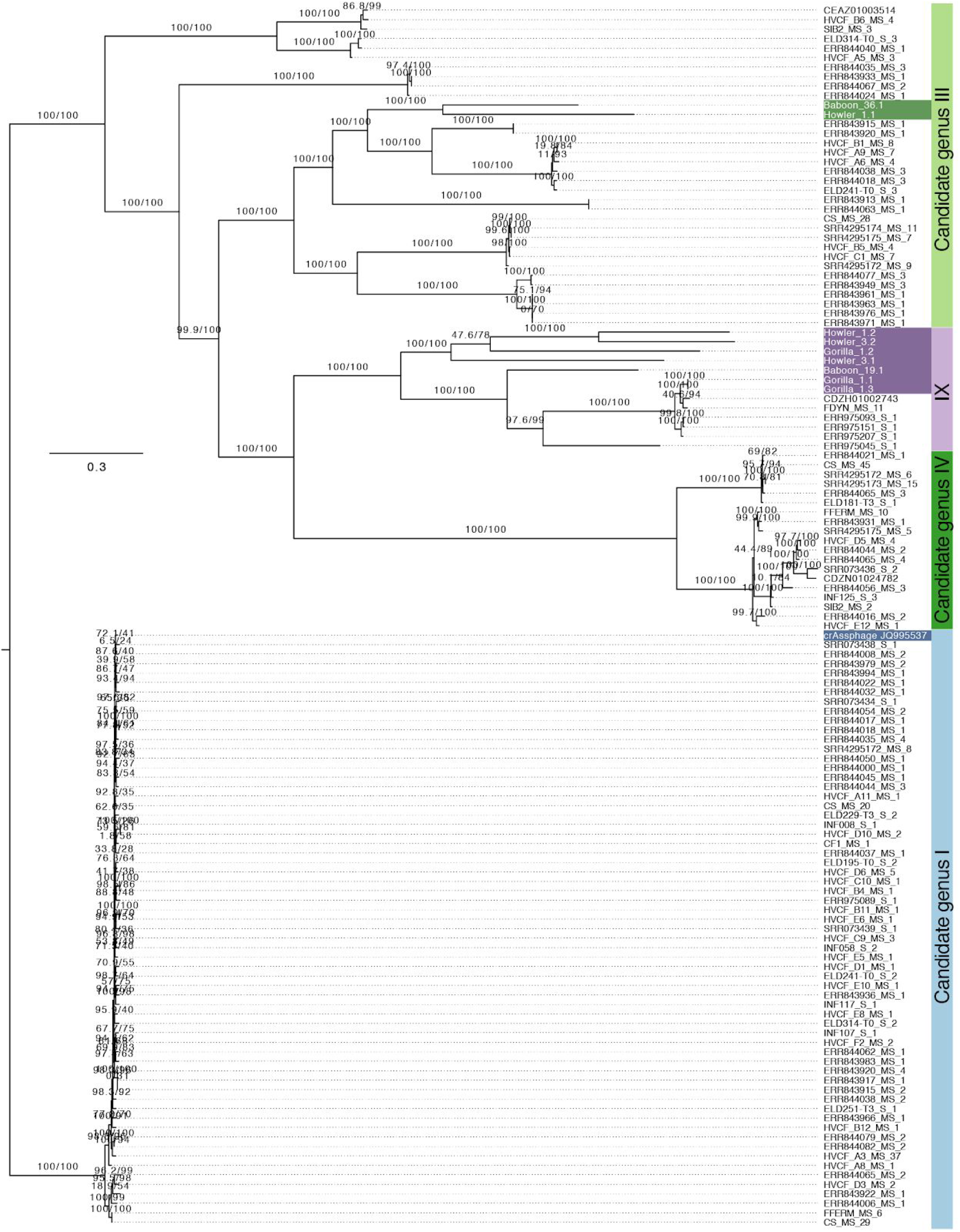
Maximum likelihood phylogeny of ten long contigs assembled from fecal metagenomes of non-human primates (highlighted) and 119 Alphacrassvirinae contigs^3^. The phylogeny is based on a 8,009 amino acid concatenated, trimmed protein alignment of 15 homologous ORFs. Branch support values are SH-aLRT and ultrafast bootstrap support^2^, scale bar indicates 0.3 mutations per site. The tree is rooted as in Guerin et al.

**Extended Data Fig. 6.**
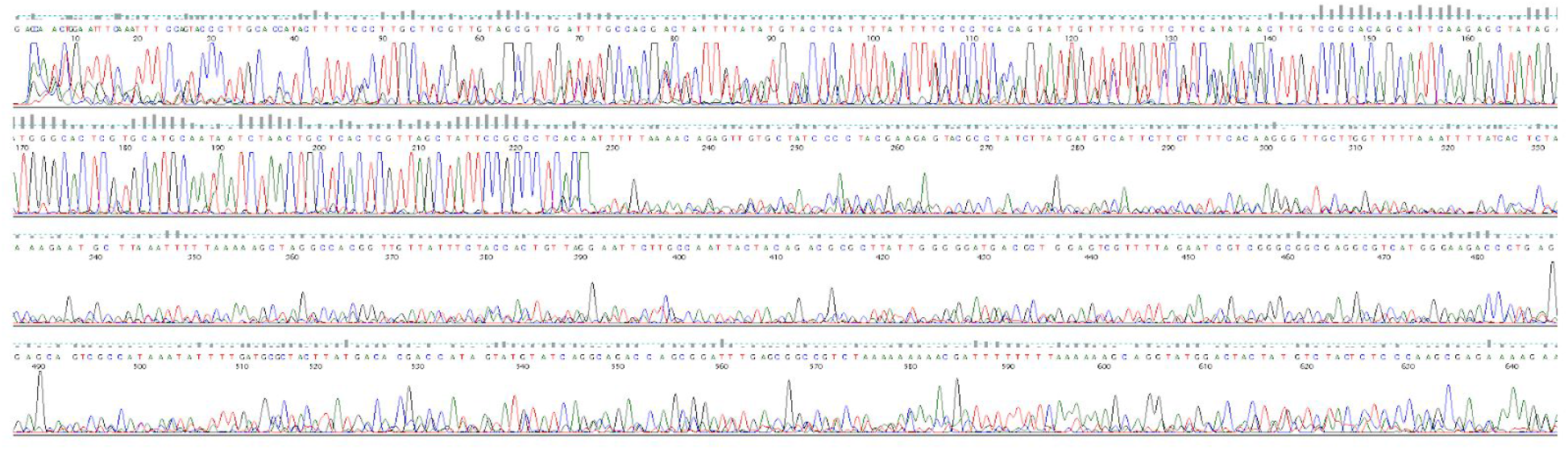
Sequencing trace of amplicon B from the wastewater treatment plant in Leuven, Belgium (sample 52GJ06_G04_B_F, see https://github.com/linsalrob/crAssohaαe/blob/master/Global_Survey/Sequences/raw_data/Lavigne/52GJ06_G04_B_F.ab1). The trace contains a single sequence for the first 227 nucleotides and then more than one sequence (presumably through an indel), rendering the trace unreadable.

**Extended Data Fig. 7.**
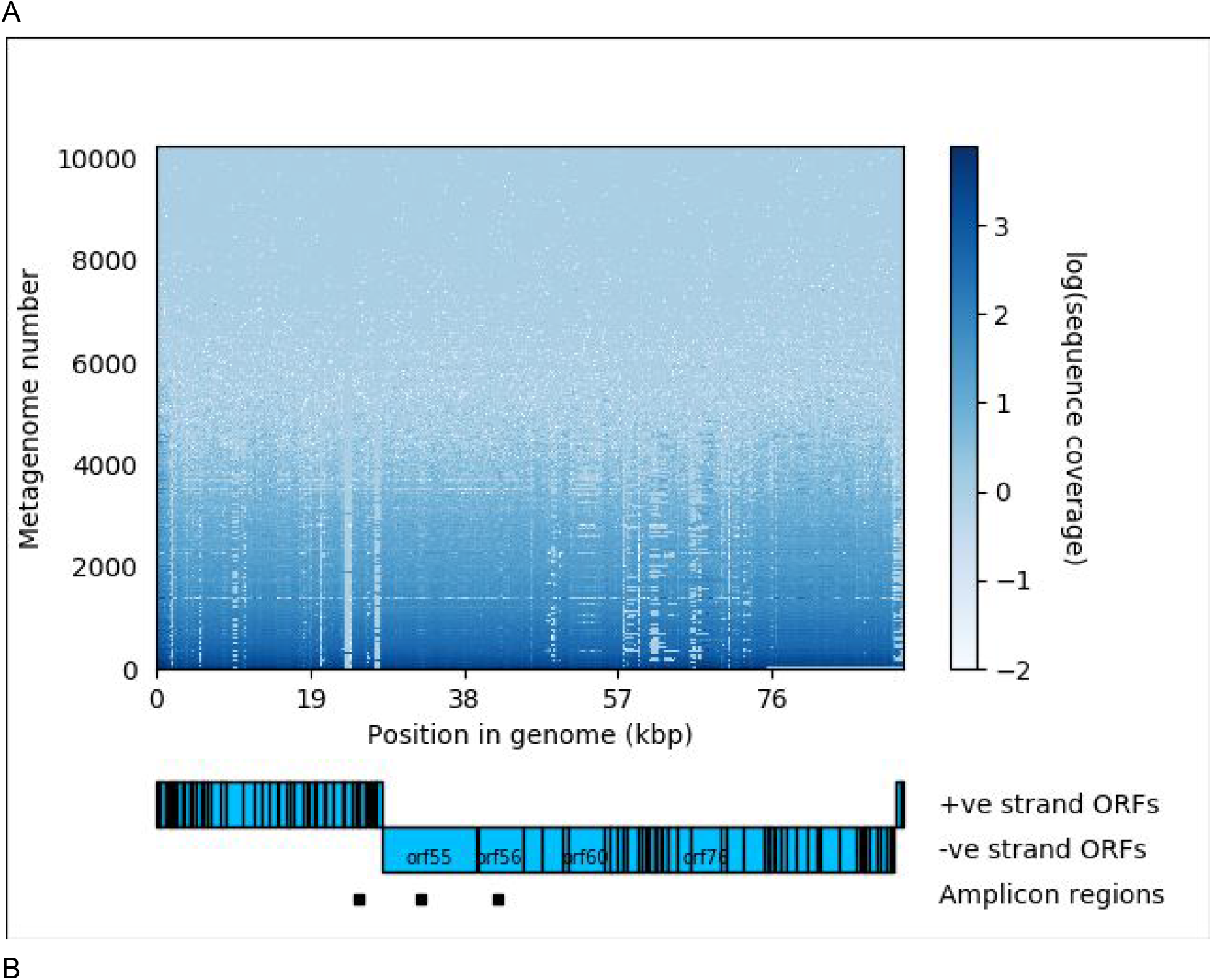
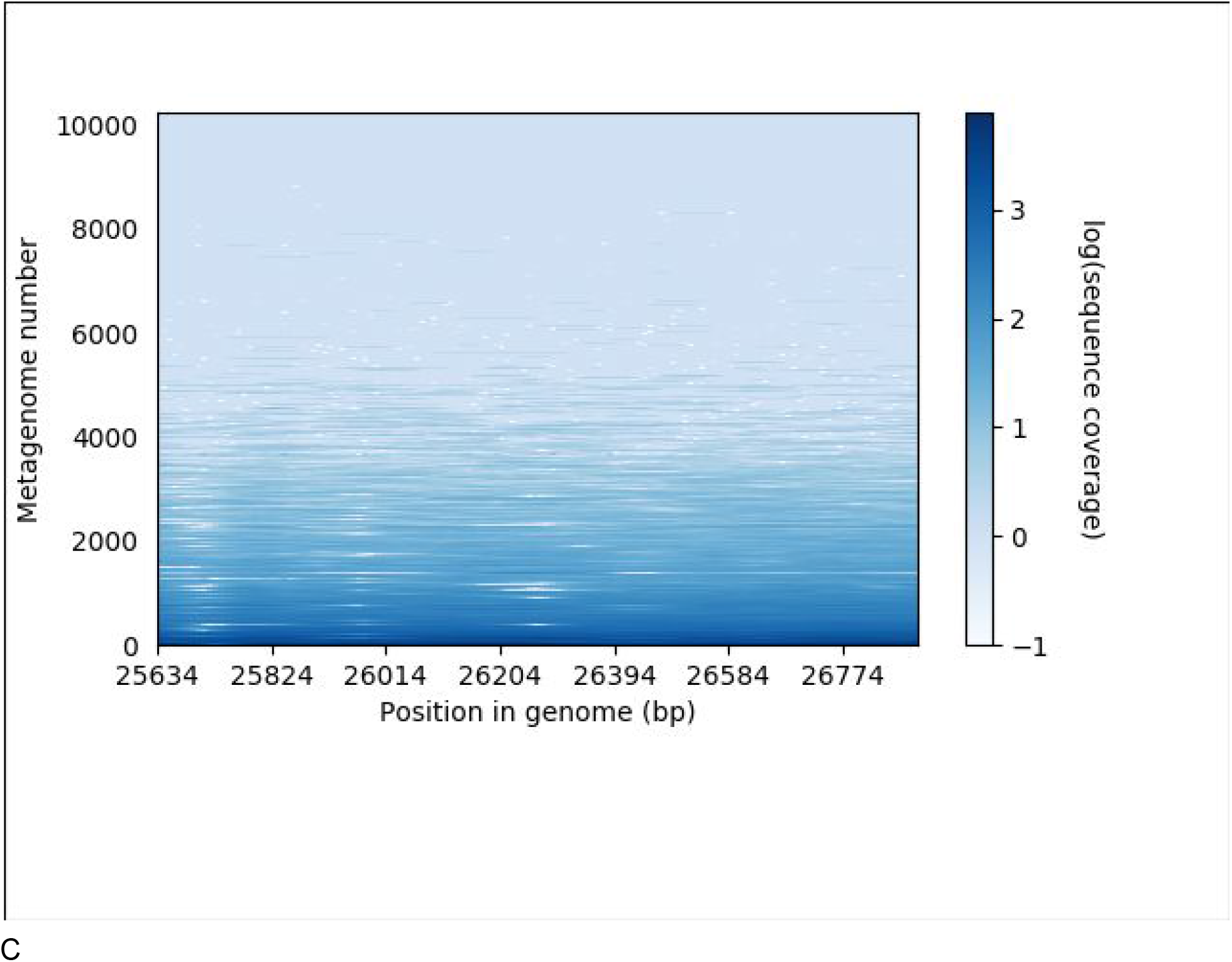
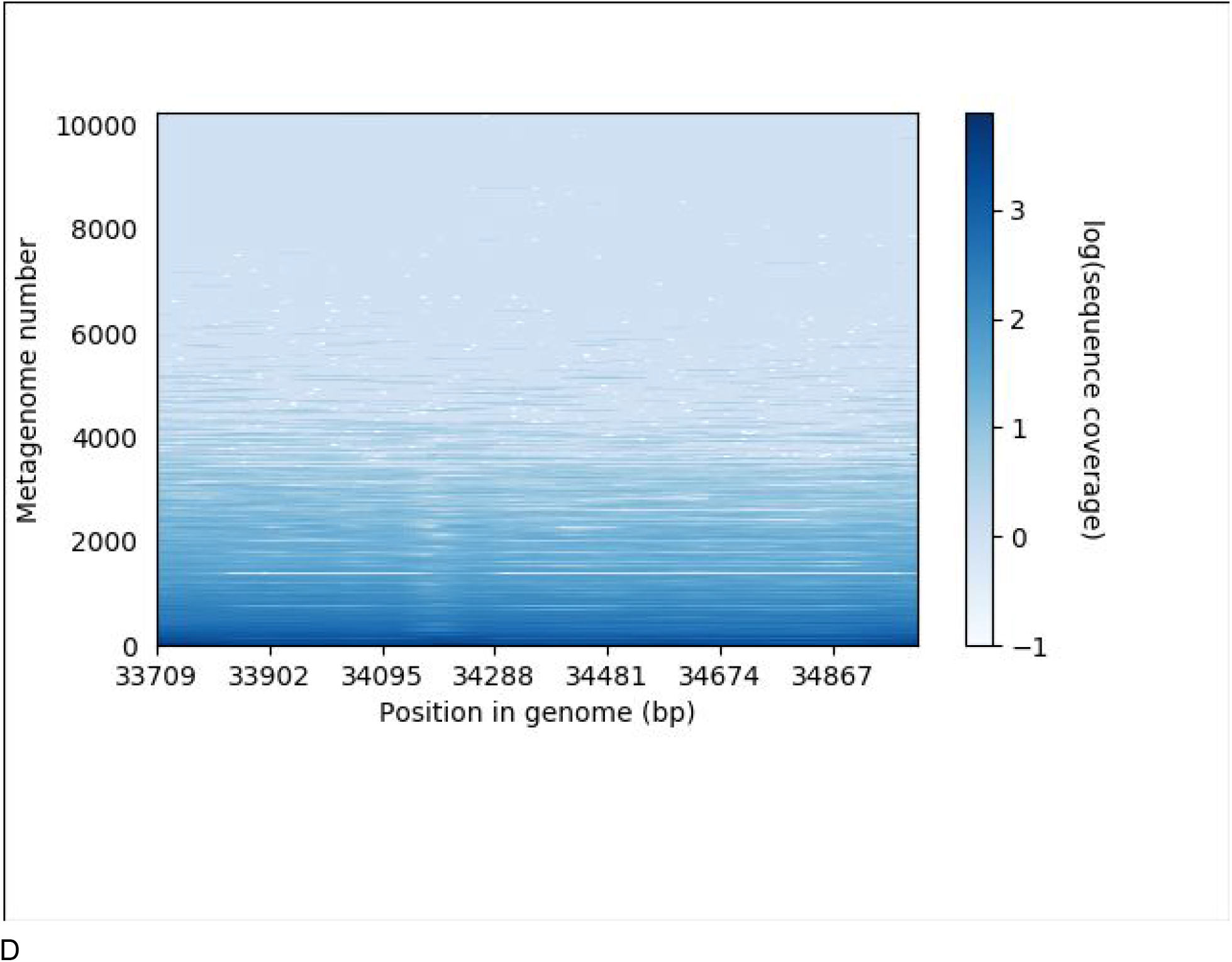
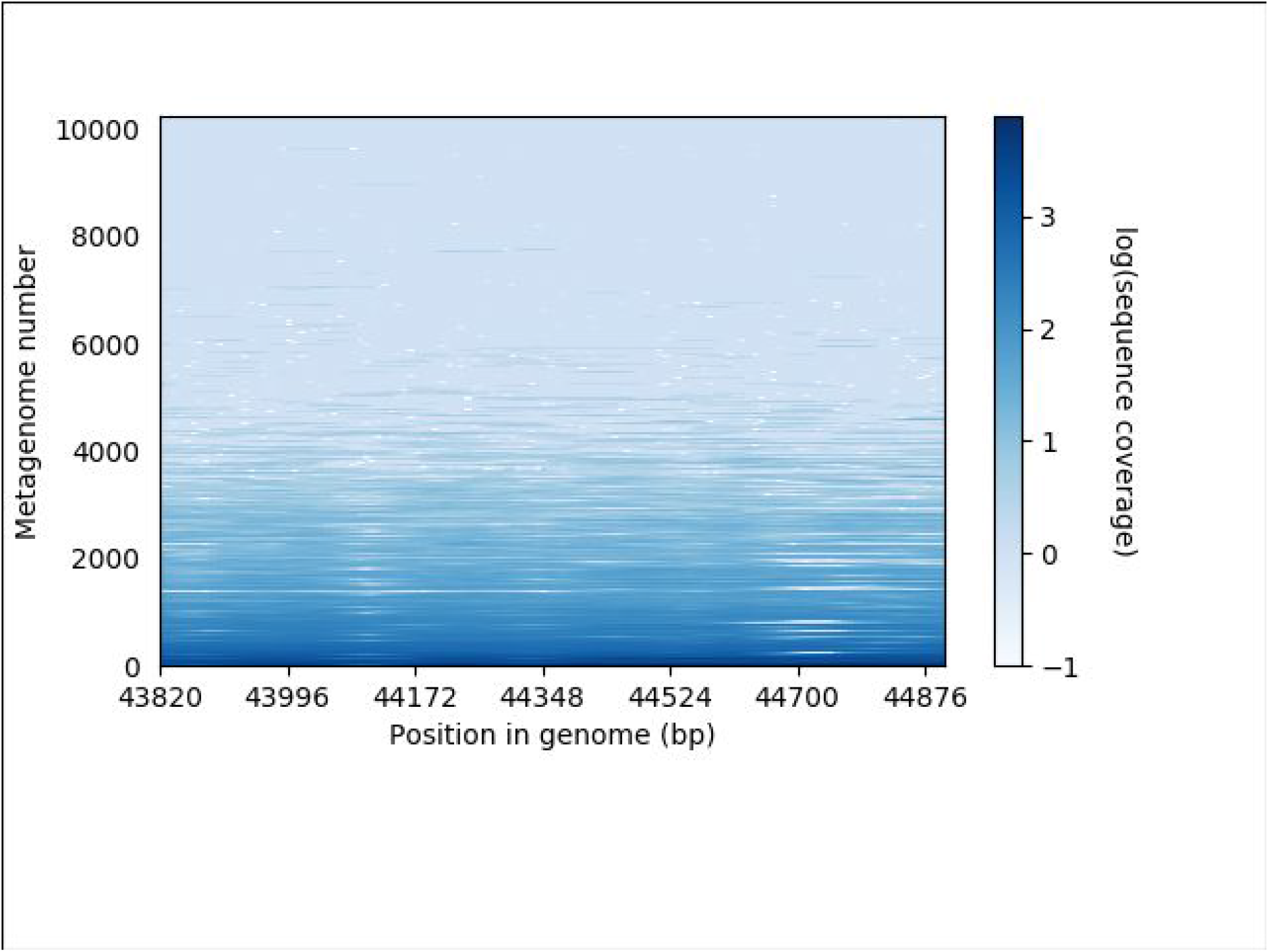
Coverage of the crAssphage genome in 10,260 metagenomes. A. Coverage across the entire genome. The predicted ORFs are shown below the genome position (x-axis) and the metagenomes are on the y-axis. Positions of the three amplicon regions are also shown. Each position represents the log of the average sequence coverage over a 1kb window as shown in the scale bar. B-D, Coverage across the three amplicon regions (A-C, respectively). Each position represents the log of the sequence coverage as shown in the scale bar.

**Extended Data Fig. 8.**
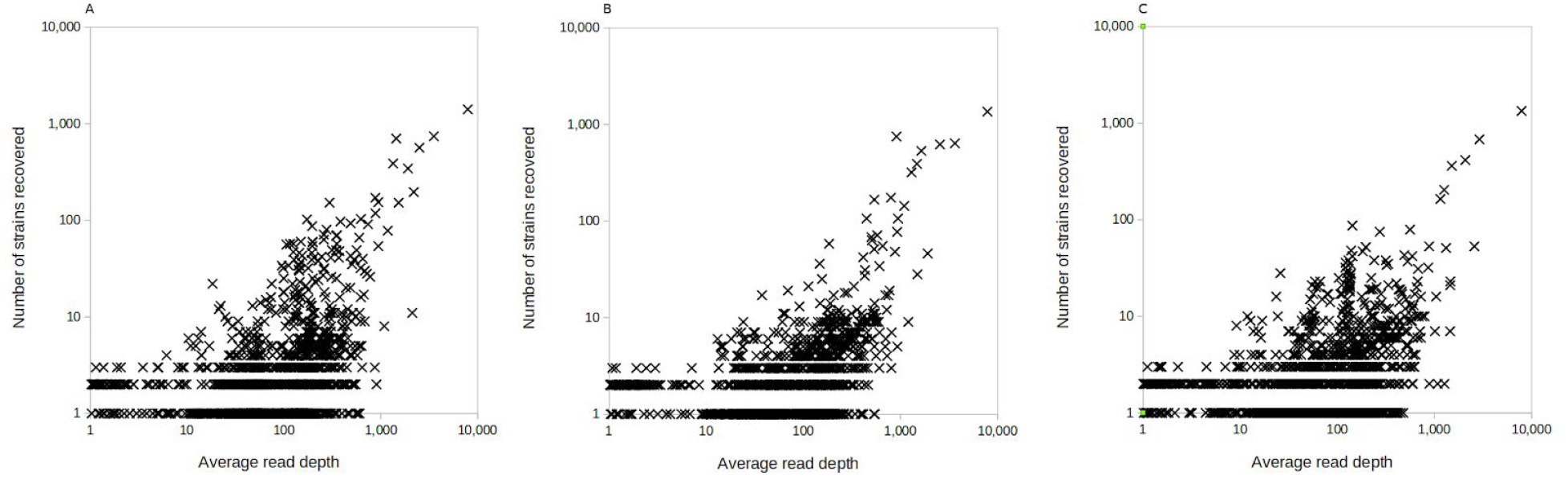
The relationship between the average per-base read depth as reported by samtools depth (including zero-coverage bases) and the number of strains recovered for amplicon A (Pearson’s ŕ=0.683; p<.001), B (Pearson’s r^2^=0.655; p<0.001), and C (Pearson’s r^2^=0.640; p<0.001).

**Extended Data Fig. 9.**
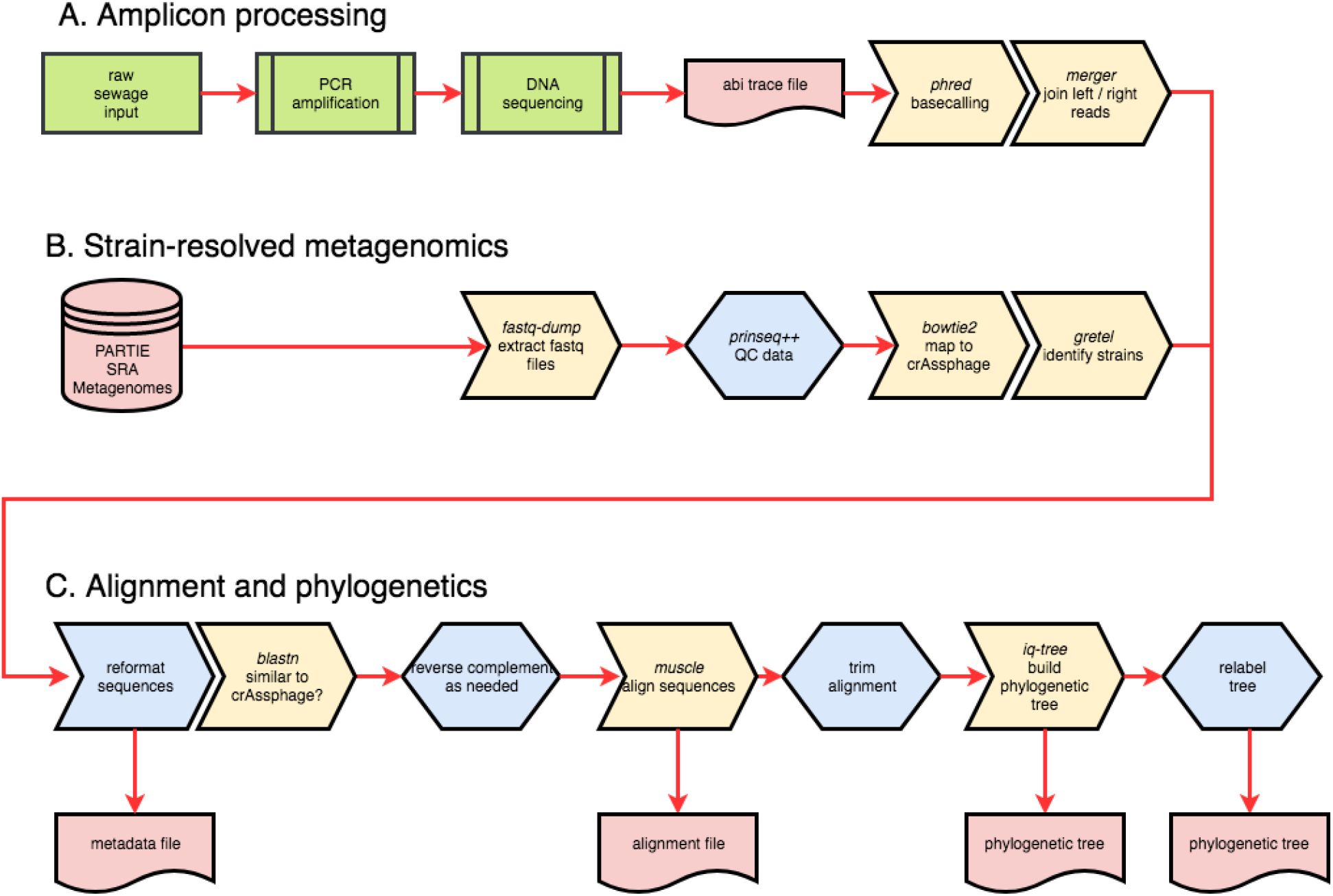
Flow chart of the sequencing analysis. Biological sample processing are shown in green, files and databases in red, external software in yellow, and software developed for this project in blue. Hexagons indicate decision steps. Amplicon sequencing starts with generating the sequences, while the metagenomics pipeline starts with publicly available sequence data. Both pipelines use the same downstream processing steps to generate the trees.

## Extended Data Tables

**Extended Data Table 1.** All crAssphage sequences collected from different sources. The numbers indicate: (i) total sequences identified, (ii) unique sequences, and (iii) sequences with locality information. The information per strain is provided in Supplementary File 2.

| Source | Amplicon A | Amplicon B | Amplicon C |
| --- | --- | --- | --- |
| Global collaboratory | 192 / 180 / 192 | 172 / 159 / 172 | 208 / 198 / 208 |
| Volunteer sequences | 27 / 27 / 27 | 11 / 11 / 11 | 25 / 25 / 25 |
| COMPARE project | 56 / 56 / 56 | 64 / 64 / 64 | 62 / 62 / 62 |
| Daycare Study | 4 / 4 / 4 | 2 / 2 / 2 | 6 / 6 / 6 |
| Strain resolved metagenomics (SRA) | 12,392 / 11,985 / 2,145 | 10,038 / 9,851 / 1,647 | 9,013 / 8,794 / 1,473 |
| Total | 12,671 / 12,252 / 2,424 | 10,287 / 10,087 / 1,896 | 9,314 / 9,083 / 1,774 |

**Extended Data Table 2.**
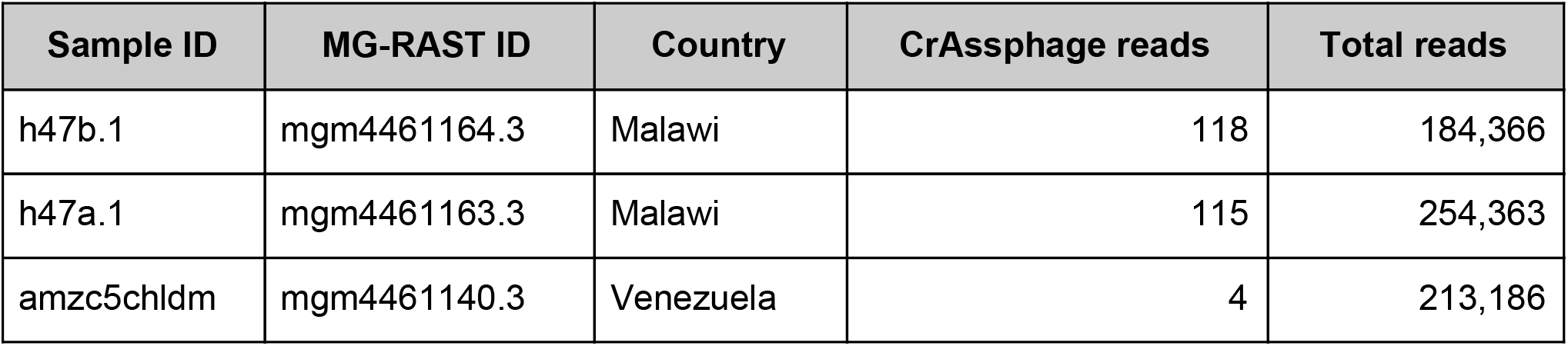
Number of crAssphage reads in fecal metagenomes from rural Malawi and the Amazonas of Venezuela^4^.

| Sample ID | MG-RAST ID | Country | CrAssphage reads | Total reads |
| --- | --- | --- | --- | --- |
| h47b.1 | mgm4461164.3 | Malawi | 118 | 184,366 |
| h47a.1 | mgm4461163.3 | Malawi | 115 | 254,363 |
| amzc5chldm | mgm4461140.3 | Venezuela | 4 | 213,186 |

**Extended Data Table 3.**
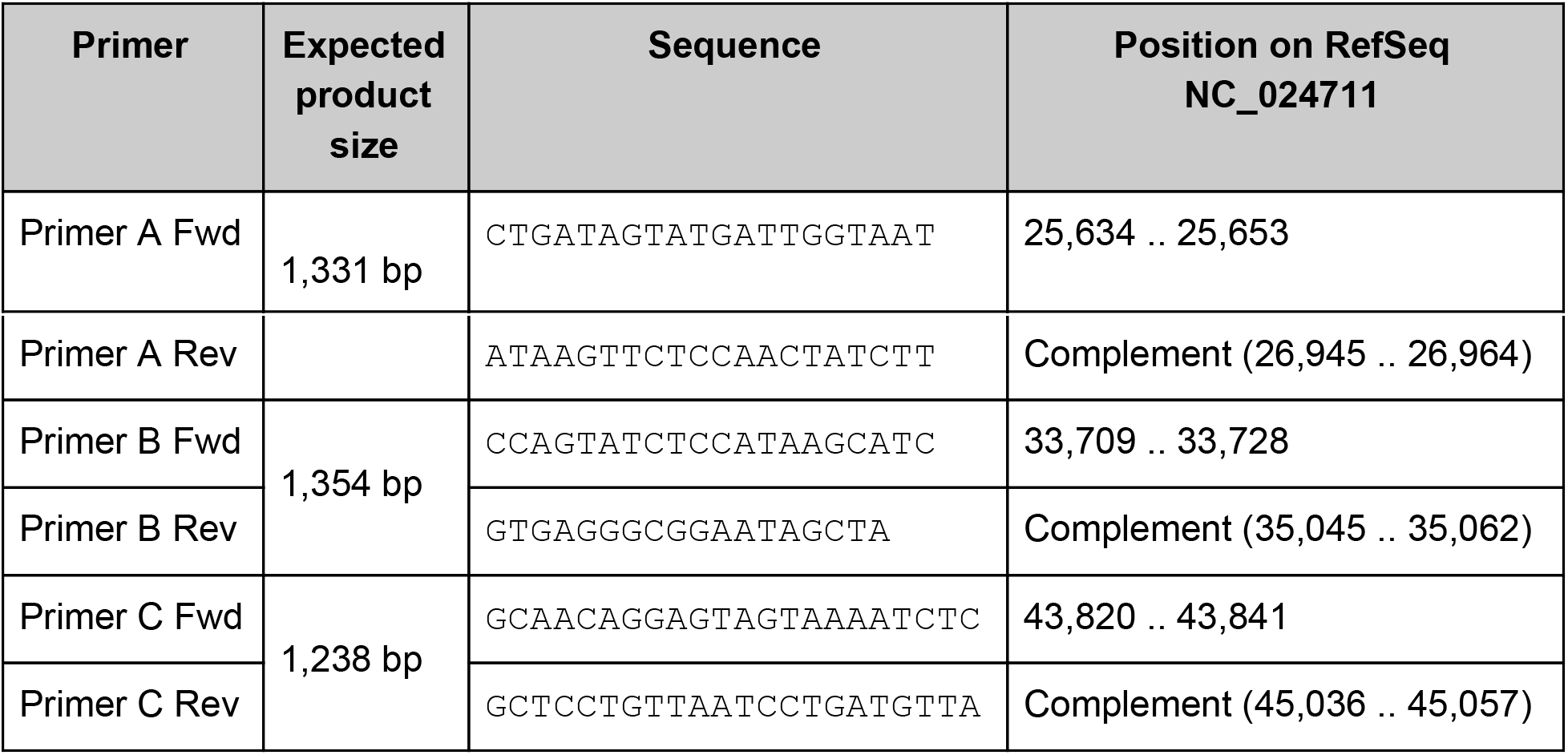
Primer sequences. Primer A, expected product size: 1,331 bp. Primer B: 1,354 bp. Primer C: 1,238 bp.

**Extended Data Table 4.** PCR reaction mixture.

| Reagent | Volume |
| --- | --- |
| DNA template | 7.0 µl |
| 2x Master Mix | 25.0 µl |
| Forward primer (10 µM) | 2.0 µl |
| Reverse primer (10 µM) | 2.0 µl |
| DNase free water | 14.0 µl |
| Total volume | 50.0 µl |

**Extended Data Table 5.** PCR amplification protocols.

| Primer A | Primers B & C |
| --- | --- |
| Denature 95°C for 3 minutes | Denature 95°C for 3 minutes |
| Then 30 cycles of:<br>Denature 95°C for 45 seconds<br>Annealing 42.6°C for 30 seconds<br>Extension 68°C for 90 seconds | Then 30 cycles of:<br>Denature 95°C for 45 seconds<br>Annealing 50°C for 30 seconds<br>Extension 68°C for 90 seconds |
| Final extension 68°C for 5 minutes | Final extension 68°C for 5 minutes |

| Name | Contents |
| --- | --- |
| Supplementary File 1 - Most ubiquitous crAssphage strain | Description and links to the bioprojects and samples that all contain identical crAssphage sequences |
| Supplementary File 2 - Global Sampling of crAssphage | The metadata and sequence data for each of the amplicon regions included in this study. |
| Supplementary File 3 - Gretel strains | Number of strains identified from all of the different samples in the SRA |
| Supplementary File 4 - Lifelines phenotype correlations | Correlation, p value, and adjusted p value for 207 exogenous and intrinsic human variables and the presence of crAssphage in stools. |
| Supplementary File 5 - Lifelines microbial correlations | Correlation, p value, and adjusted p value for the presence of 491 bacterial isolates and the presence of crAssphage in stools. |

