## Supplementary material for "Global phylogeography and ancient evolution of the widespread human gut virus crAssphage": Most ubiquitous crAssphage strain

**Supplemental File 1. The 104 samples containing the most ubiquitous crAssphage strain.**

| **Project** | **Title** | **Samples** | **Location** | **Lat/Lon** |
| --- | --- | --- | --- | --- |
|  | Seasonal Dynamics of DNA and RNA Viral Bioaerosol Communities in a Daycare Setting | Daycare study sample AP-DNA-10 | Virginia, USA |  |
| [ERP008729](https://trace.ncbi.nlm.nih.gov/Traces/sra/?study=ERP008729) | Gut microbiome development along the colorectal adenoma-carcinoma sequence | [ERR688473](https://www.ncbi.nlm.nih.gov/sra/ERR688473) |  |  |
| [SRP059928](https://trace.ncbi.nlm.nih.gov/Traces/sra/?study=SRP059928) | Non-human sequence from stool, colon biopsy, ileum resection, kefir, and artificial bacterial mixtures | [SRR2082443](https://www.ncbi.nlm.nih.gov/sra/SRR2082443) | Canada | 53.520710 N 113.524239 W |
| [SRP065270](https://trace.ncbi.nlm.nih.gov/Traces/sra/?study=SRP065270) | Functional dynamics of the elderly gut microbiome during probiotic consumption | [SRR2857970](https://www.ncbi.nlm.nih.gov/sra/SRR2857970) | USA | 42.3601 N 71.0589 W |
| [ERP002061](https://trace.ncbi.nlm.nih.gov/Traces/sra/?study=ERP002061) | A method for identifying metagenomic species and variable genetic elements by exhaustive co-abundance binning | [ERR210123](https://www.ncbi.nlm.nih.gov/sra/ERR210123); [ERR210122](https://www.ncbi.nlm.nih.gov/sra/ERR210122); [ERR210052](https://www.ncbi.nlm.nih.gov/sra/ERR210052); [ERR209575](https://www.ncbi.nlm.nih.gov/sra/ERR209575); [ERR209644](https://www.ncbi.nlm.nih.gov/sra/ERR209644) |  |  |
| [ERP005860](https://trace.ncbi.nlm.nih.gov/Traces/sra/?study=ERP005860) | Liver cirrhosis occurs as a consequence of many chronic liver diseases that are prevalent worldwide. Previous studies have shown an association between the gut microbiota and liver complications such as cirrhosis and other liver injuries. We therefore undertook a whole gut microbiome wide association study of stool samples from 98 liver cirrhosis patients and 83 healthy controls to characterise the faecal microbial communities and their functional composition. In total, we generated 860 Gb of high-quality sequence data and built a reference gene set for the liver cirrhosis cohort containing 2.69 million genes, 36.1% of which was not covered by previously published gene catalogues. | [ERR527052](https://www.ncbi.nlm.nih.gov/sra/ERR527052) |  |  |
| [SRP083099](https://trace.ncbi.nlm.nih.gov/Traces/sra/?study=SRP083099) | Gut microbiota and metagenomic diversity of omnivore, vegetarian and vegan healthy subjects | [SRR4074354](https://www.ncbi.nlm.nih.gov/sra/SRR4074354) | Italy |  |
| [ERP000108](https://trace.ncbi.nlm.nih.gov/Traces/sra/?study=ERP000108) | A human gut microbial gene catalog established by deep metagenomic sequencing | [ERR011190](https://www.ncbi.nlm.nih.gov/sra/ERR011190) |  |  |
| [DRP000700](https://trace.ncbi.nlm.nih.gov/Traces/sra/?study=DRP000700) | Metgenomic analysis of human gut microbiome in patients with multiple sclerosis (MS). | [DRR002666](https://www.ncbi.nlm.nih.gov/sra/DRR002666) |  |  |
| [SRP056641](https://trace.ncbi.nlm.nih.gov/Traces/sra/?study=SRP056641) | Human Microbiome Environment Metagenome | [SRR2175726](https://www.ncbi.nlm.nih.gov/sra/SRR2175726) |  |  |
| [SRP066479](https://trace.ncbi.nlm.nih.gov/Traces/sra/?study=SRP066479) | Antibiotic resistance exchange between microbiota in resource-poor settings in Latin America | [SRR2938428](https://www.ncbi.nlm.nih.gov/sra/SRR2938428) | El Salvador |  |
| [ERP005534](https://trace.ncbi.nlm.nih.gov/Traces/sra/?study=ERP005534) | Potential of fecal microbiota for early stage detection of colorectal cancer | [ERR480821](https://www.ncbi.nlm.nih.gov/sra/ERR480821); [ERR479008](https://www.ncbi.nlm.nih.gov/sra/ERR479008); [ERR480516](https://www.ncbi.nlm.nih.gov/sra/ERR480516); [ERR479525](https://www.ncbi.nlm.nih.gov/sra/ERR479525); [ERR480673](https://www.ncbi.nlm.nih.gov/sra/ERR480673); [ERR479524](https://www.ncbi.nlm.nih.gov/sra/ERR479524); [ERR480711](https://www.ncbi.nlm.nih.gov/sra/ERR480711) |  |  |
| [SRP029441](https://trace.ncbi.nlm.nih.gov/Traces/sra/?study=SRP029441) | Fiji COMP | [SRR2195841](https://www.ncbi.nlm.nih.gov/sra/SRR2195841); [SRR2249814](https://www.ncbi.nlm.nih.gov/sra/SRR2249814); [SRR2222814](https://www.ncbi.nlm.nih.gov/sra/SRR2222814); [SRR2250644](https://www.ncbi.nlm.nih.gov/sra/SRR2250644); [SRR2189708](https://www.ncbi.nlm.nih.gov/sra/SRR2189708) |  |  |
| [SRP049045](https://trace.ncbi.nlm.nih.gov/Traces/sra/?study=SRP049045) | Abundance of antibiotic resistance genes and structure of the microbial community in wastewater of medical facilities besides hospitals | [SRR1616987](https://www.ncbi.nlm.nih.gov/sra/SRR1616987) | Germany | 48 N 8 E |
| [SRP072561](https://trace.ncbi.nlm.nih.gov/Traces/sra/?study=SRP072561) | Human Gut Microbiome in a Multiplex Family Study of Type 1 Diabetes Mellitus | [SRR3313047](https://www.ncbi.nlm.nih.gov/sra/SRR3313047) | Luxembourg | 49.5 N 6.2 E |
| [ERP013562](https://trace.ncbi.nlm.nih.gov/Traces/sra/?study=ERP013562) | Gut microbial dysbiosis in young adults with obesity | [ERR1190645](https://www.ncbi.nlm.nih.gov/sra/ERR1190645); [ERR1190689](https://www.ncbi.nlm.nih.gov/sra/ERR1190689); [ERR1190633](https://www.ncbi.nlm.nih.gov/sra/ERR1190633) |  |  |
| [SRP060568](https://trace.ncbi.nlm.nih.gov/Traces/sra/?study=SRP060568) | Hospital Air Samples Metagenome | [SRR2183670](https://www.ncbi.nlm.nih.gov/sra/SRR2183670) |  |  |
| [SRP066514](https://trace.ncbi.nlm.nih.gov/Traces/sra/?study=SRP066514) | Human gut Metagenome | [SRR2940957](https://www.ncbi.nlm.nih.gov/sra/SRR2940957) | USA |  |
| [ERP013933](https://trace.ncbi.nlm.nih.gov/Traces/sra/?study=ERP013933) | Reproducibility of associations between the human gut microbiome and colorectal cancer assessed in a patient population from Washington, DC, USA | [ERR1293299](https://www.ncbi.nlm.nih.gov/sra/ERR1293299); [ERR1293522](https://www.ncbi.nlm.nih.gov/sra/ERR1293522) |  |  |
| [ERP012929](https://trace.ncbi.nlm.nih.gov/Traces/sra/?study=ERP012929) | Towards personalized nutrition by prediction of glycemic responses | [ERR1137395](https://www.ncbi.nlm.nih.gov/sra/ERR1137395); [ERR1137041](https://www.ncbi.nlm.nih.gov/sra/ERR1137041); [ERR1136988](https://www.ncbi.nlm.nih.gov/sra/ERR1136988) |  |  |
| [DRP000446](https://trace.ncbi.nlm.nih.gov/Traces/sra/?study=DRP000446) | Comprehensive Detection of Possible Pathogens Associated with Kawasaki Disease | [DRR014146](https://www.ncbi.nlm.nih.gov/sra/DRR014146) |  |  |
| [SRP002163](https://trace.ncbi.nlm.nih.gov/Traces/sra/?study=SRP002163) | Human Microbiome Project (HMP) Metagenomic WGS Projects, deeper sequencing of the human microbiome samples: Production Phase | [SRR528262](https://www.ncbi.nlm.nih.gov/sra/SRR528262); [SRR1804206](https://www.ncbi.nlm.nih.gov/sra/SRR1804206); [SRR532466](https://www.ncbi.nlm.nih.gov/sra/SRR532466); [SRR1804707](https://www.ncbi.nlm.nih.gov/sra/SRR1804707); [SRR532351](https://www.ncbi.nlm.nih.gov/sra/SRR532351); [SRR514308](https://www.ncbi.nlm.nih.gov/sra/SRR514308); [SRR063906](https://www.ncbi.nlm.nih.gov/sra/SRR063906); [SRR539598](https://www.ncbi.nlm.nih.gov/sra/SRR539598); [SRR1804115](https://www.ncbi.nlm.nih.gov/sra/SRR1804115); [SRR549428](https://www.ncbi.nlm.nih.gov/sra/SRR549428); [SRR532027](https://www.ncbi.nlm.nih.gov/sra/SRR532027) |  |  |
| [SRP051174](https://trace.ncbi.nlm.nih.gov/Traces/sra/?study=SRP051174) | NIBSC_BSRI Metagenome | [SRR1714192](https://www.ncbi.nlm.nih.gov/sra/SRR1714192) | USA | 38.98 N 77.11 W |
| [SRP064913](https://trace.ncbi.nlm.nih.gov/Traces/sra/?study=SRP064913) | Library preparation methodology can influence genomic and functional predictions in human microbiome research | [SRR2726666](https://www.ncbi.nlm.nih.gov/sra/SRR2726666) |  |  |
| [ERP009422](https://trace.ncbi.nlm.nih.gov/Traces/sra/?study=ERP009422) | Temporal and technical variability of human gut metagenomes | [ERR748434](https://www.ncbi.nlm.nih.gov/sra/ERR748434); [ERR748433](https://www.ncbi.nlm.nih.gov/sra/ERR748433); [ERR748174](https://www.ncbi.nlm.nih.gov/sra/ERR748174); [ERR748184](https://www.ncbi.nlm.nih.gov/sra/ERR748184); [ERR748319](https://www.ncbi.nlm.nih.gov/sra/ERR748319); [ERR748477](https://www.ncbi.nlm.nih.gov/sra/ERR748477) |  |  |
| [SRP000319](https://trace.ncbi.nlm.nih.gov/Traces/sra/?study=SRP000319) | A core gut microbiome in obese and lean twins | [SRR029696](https://www.ncbi.nlm.nih.gov/sra/SRR029696) |  |  |
| [SRP058816](https://trace.ncbi.nlm.nih.gov/Traces/sra/?study=SRP058816) | Methanogenic digester communities Raw sequence reads | [SRR2043640](https://www.ncbi.nlm.nih.gov/sra/SRR2043640) |  |  |
| [SRP040146](https://trace.ncbi.nlm.nih.gov/Traces/sra/?study=SRP040146) | C.diff FMT | [SRR1492958](https://www.ncbi.nlm.nih.gov/sra/SRR1492958); [SRR1437940](https://www.ncbi.nlm.nih.gov/sra/SRR1437940); [SRR1491454](https://www.ncbi.nlm.nih.gov/sra/SRR1491454); [SRR1437798](https://www.ncbi.nlm.nih.gov/sra/SRR1437798); [SRR1462693](https://www.ncbi.nlm.nih.gov/sra/SRR1462693); [SRR1437716](https://www.ncbi.nlm.nih.gov/sra/SRR1437716); [SRR1437790](https://www.ncbi.nlm.nih.gov/sra/SRR1437790); [SRR1461800](https://www.ncbi.nlm.nih.gov/sra/SRR1461800); [SRR1491724](https://www.ncbi.nlm.nih.gov/sra/SRR1491724); [SRR1490908](https://www.ncbi.nlm.nih.gov/sra/SRR1490908); [SRR1461818](https://www.ncbi.nlm.nih.gov/sra/SRR1461818); [SRR1490972](https://www.ncbi.nlm.nih.gov/sra/SRR1490972); [SRR1490923](https://www.ncbi.nlm.nih.gov/sra/SRR1490923) |  |  |
| [ERP013563](https://trace.ncbi.nlm.nih.gov/Traces/sra/?study=ERP013563) | Gut microbiome-dependent stratification of patients for anti-diabetic treatment | [ERR1190879](https://www.ncbi.nlm.nih.gov/sra/ERR1190879); [ERR1190804](https://www.ncbi.nlm.nih.gov/sra/ERR1190804) |  |  |
| [SRP056054](https://trace.ncbi.nlm.nih.gov/Traces/sra/?study=SRP056054) | A prospective, longitudinal analysis of the developing gut microbiome in infants en route to type 1 diabetes | [SRR1918833](https://www.ncbi.nlm.nih.gov/sra/SRR1918833); [SRR1910622](https://www.ncbi.nlm.nih.gov/sra/SRR1910622) |  |  |
| [SRP031463](https://trace.ncbi.nlm.nih.gov/Traces/sra/?study=SRP031463) | Microbiome analysis of stool samples from African Americans with colon polyps | [SRR1012404](https://www.ncbi.nlm.nih.gov/sra/SRR1012404) |  |  |
| [SRP040765](https://trace.ncbi.nlm.nih.gov/Traces/sra/?study=SRP040765) | Microbiome study of the RISK cohort | [SRR1765589](https://www.ncbi.nlm.nih.gov/sra/SRR1765589) |  |  |
| [SRP064400](https://trace.ncbi.nlm.nih.gov/Traces/sra/?study=SRP064400) | Intestinal microbiota dynamics in hospitalized patients | [SRR2565987](https://www.ncbi.nlm.nih.gov/sra/SRR2565987); [SRR2565536](https://www.ncbi.nlm.nih.gov/sra/SRR2565536); [SRR2566055](https://www.ncbi.nlm.nih.gov/sra/SRR2566055) | Canada | 45.50 N 73.63 W |
| [SRP011011](https://trace.ncbi.nlm.nih.gov/Traces/sra/?study=SRP011011) | A Metagenome-Wide Association Study of gut microbiota identifies markers associated with Type 2 Diabetes | [SRR413683](https://www.ncbi.nlm.nih.gov/sra/SRR413683) |  |  |
| [ERP016813](https://trace.ncbi.nlm.nih.gov/Traces/sra/?study=ERP016813) | Integrated metabolomics and metagenomics analysis of plasma and urine identified microbial metabolites associated with coronary heart disease | [ERR1578695](https://www.ncbi.nlm.nih.gov/sra/ERR1578695) |  |  |
| [ERP009131](https://trace.ncbi.nlm.nih.gov/Traces/sra/?study=ERP009131) | The initial state of the human gut microbiome determines its reshaping by antibiotics | [ERR719489](https://www.ncbi.nlm.nih.gov/sra/ERR719489); [ERR719406](https://www.ncbi.nlm.nih.gov/sra/ERR719406); [ERR719401](https://www.ncbi.nlm.nih.gov/sra/ERR719401); [ERR719424](https://www.ncbi.nlm.nih.gov/sra/ERR719424); [ERR719642](https://www.ncbi.nlm.nih.gov/sra/ERR719642) |  |  |
| [ERP003612](https://trace.ncbi.nlm.nih.gov/Traces/sra/?study=ERP003612) | Richness of human gut microbiome correlates with metabolic markers | [ERR321165](https://www.ncbi.nlm.nih.gov/sra/ERR321165) |  |  |
| [ERP005989](https://trace.ncbi.nlm.nih.gov/Traces/sra/?study=ERP005989) | Dynamics and Stabilization of the Human Gut Microbiome during the First Year of Life | [ERR525816](https://www.ncbi.nlm.nih.gov/sra/ERR525816) |  |  |
| [SRP080787](https://trace.ncbi.nlm.nih.gov/Traces/sra/?study=SRP080787) | Mongolian Metagenome | [SRR3992959](https://www.ncbi.nlm.nih.gov/sra/SRR3992959) | China | 43.95 N 116.16 E |
| [SRP059392](https://trace.ncbi.nlm.nih.gov/Traces/sra/?study=SRP059392) | Ecological reactor Metagenome | [SRR2062623](https://www.ncbi.nlm.nih.gov/sra/SRR2062623) | USA | 43.727094 N 72.425964 W |
| [SRP008047](https://trace.ncbi.nlm.nih.gov/Traces/sra/?study=SRP008047) | A Metagenome-Wide Association Study of gut microbiota identifies markers associated with Type 2 Diabetes | [SRR341594](https://www.ncbi.nlm.nih.gov/sra/SRR341594) |  |  |
| [DRP003048](https://trace.ncbi.nlm.nih.gov/Traces/sra/?study=DRP003048) | Metagenomics of Japanese gut microbiomes | [DRR042632](https://www.ncbi.nlm.nih.gov/sra/DRR042632); [DRR042593](https://www.ncbi.nlm.nih.gov/sra/DRR042593); [DRR042410](https://www.ncbi.nlm.nih.gov/sra/DRR042410) |  |  |
| [SRP002523](https://trace.ncbi.nlm.nih.gov/Traces/sra/?study=SRP002523) | Metagenomic analysis of viruses in the fecal micorobiota of monozygotic twins and their mothers | [SRR073436](https://www.ncbi.nlm.nih.gov/sra/SRR073436); [SRR073432](https://www.ncbi.nlm.nih.gov/sra/SRR073432) |  |  |
| [ERP003671](https://trace.ncbi.nlm.nih.gov/Traces/sra/?study=ERP003671) | Deep Illumina-based shotgun sequencing reveals dietary effects on the structure and function of the faecal microbiome of growing kittens | [ERR318688](https://www.ncbi.nlm.nih.gov/sra/ERR318688) | USA |  |
| [ERP014654](https://trace.ncbi.nlm.nih.gov/Traces/sra/?study=ERP014654) | microbial diversity and function | [ERR1333182](https://www.ncbi.nlm.nih.gov/sra/ERR1333182) |  |  |
| [ERP004605](https://trace.ncbi.nlm.nih.gov/Traces/sra/?study=ERP004605) | An integrated catalog of reference genes in the human gut microbiome | [ERR414735](https://www.ncbi.nlm.nih.gov/sra/ERR414735); [ERR414539](https://www.ncbi.nlm.nih.gov/sra/ERR414539) | Spain:Madrid | 40.463667,-3.74922 |

Notes:

1. The sample from the daycare study is not yet available from the SRA.
2. Location, latitude, and longitude are provided when they are known.
